# Synthetic G protein-coupled receptors for programmable sensing and control of cell behavior

**DOI:** 10.1101/2024.04.15.589622

**Authors:** Nicholas A. Kalogriopoulos, Reika Tei, Yuqi Yan, Matthew Ravalin, Yulong Li, Alice Ting

## Abstract

Synthetic receptors that mediate antigen-dependent cell responses are transforming therapeutics, drug discovery, and basic research. However, established technologies such as chimeric antigen receptors (CARs) can only detect immobilized antigens, have limited output scope, and lack built-in drug control. Here, we engineer synthetic G protein-coupled receptors (GPCRs) capable of driving a wide range of native or nonnative cellular processes in response to user-defined antigen. We achieve modular antigen gating by engineering and fusing a conditional auto-inhibitory domain onto GPCR scaffolds. Antigen binding to a fused nanobody relieves auto-inhibition and enables receptor activation by drug, thus generating Programmable Antigen-gated G protein-coupled Engineered Receptors (PAGERs). We create PAGERs responsive to more than a dozen biologically and therapeutically important soluble and cell surface antigens, in a single step, from corresponding nanobody binders. Different PAGER scaffolds permit antigen binding to drive transgene expression, real-time fluorescence, or endogenous G protein activation, enabling control of cytosolic Ca^2+^, lipid signaling, cAMP, and neuronal activity. Due to its modular design and generalizability, we expect PAGER to have broad utility in discovery and translational science.

## Main

Cell surface receptors sense specific extracellular cues, transmit those signals across the cell membrane, and convert them into defined cellular responses. Engineering modular synthetic receptors capable of recapitulating this transmembrane signaling is a key challenge for reprogramming cell behavior. Synthetic receptors derived from T cell receptors (CARs^1,2^) and the Notch receptor (synNotch^3^) have enabled diverse applications in medicine^4,5^ and basic research.^6,7^ However, these platforms are limited by their inherent mechanisms of activation (antigen-induced clustering and force, respectively) which restrict both antigen and output scope. G protein-coupled receptors (GPCRs), the largest and most diverse family of cell surface receptors, would offer a more flexible scaffold for programming cellular responses with synthetic receptors. GPCRs are seven-transmembrane cell surface receptors that mediate responses to diverse extracellular signals, including hormones, neurotransmitters, peptides, light, force, and odorants. Ligand binding induces a conformational change in the GPCR that, in turn, activates heterotrimeric G proteins and downstream intracellular signaling cascades.

The challenge in harnessing GPCRs for synthetic receptor technology is that they are not structurally modular proteins. Previous efforts to alter ligand specificity have required labor-intensive structure-guided mutagenesis and directed evolution^8,9^. To enable straightforward and modular antigen gating, we engineered a conditional auto-inhibitory domain for GPCR scaffolds. We fused a nanobody and a receptor auto-inhibitory domain to the extracellular N-terminus of the GPCR such that binding of the auto-inhibitory domain to the GPCR and binding of the nanobody to an antigen are mutually exclusive. Thereby, antigen binding relieves auto-inhibition and enables receptor activation by an agonist (**Fig. 1a**). PAGER activation can then drive diverse native or synthetic outputs, including transgene expression via SPARK^10–13^ (PAGER_TF_), endogenous G protein activation (PAGER_G_), or real-time fluorescence via GRABs^14,15^ (PAGER_FL_).

**Figure 1.**
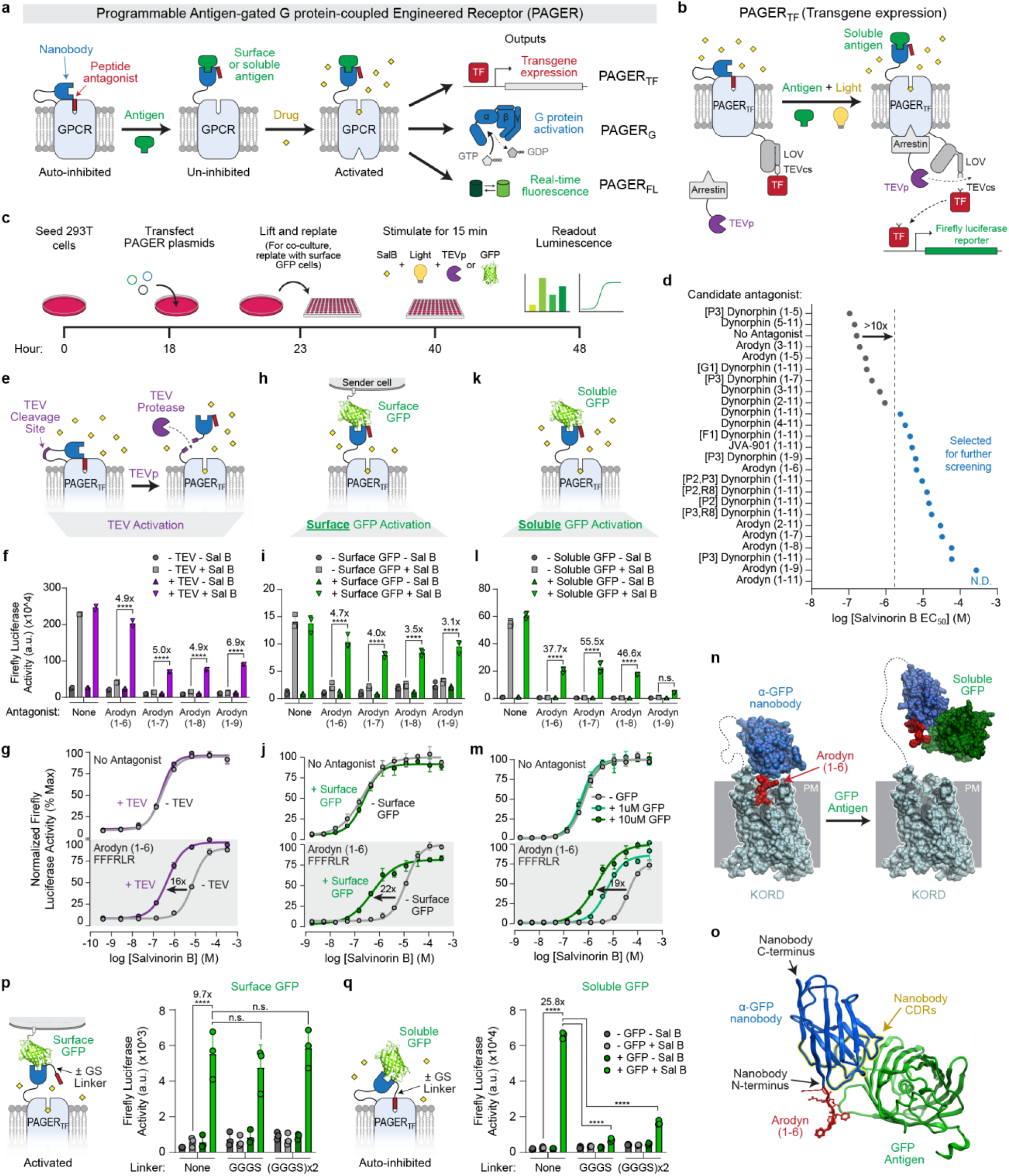
Design and optimization of PAGER. **a,** PAGER overview. A drug-activated GPCR is auto-inhibited by an N-terminally fused antagonist peptide (red). A nanobody binder (blue) is inserted next to the antagonist peptide such that binding of extracellular target antigen (green) sterically interferes with the antagonist peptide and relieves autoinhibition, enabling drug (yellow) to activate the GPCR. PAGER can respond to both cell-surface and soluble antigens and give a variety of intracellular outputs, including transgene expression, G protein activation, and real-time fluorescence. **b,** For PAGER-driven transgene expression (PAGER_TF_), the intracellular C-terminus is fused to a transcription factor (TF) via a protease-sensitive linker and LOV domain^10,11^. Upon PAGER activation, arrestin-TEV protease (TEVp) is recruited to the receptor, leading to proximity-dependent release of the TF and reporter gene expression (FLuc, firefly luciferase). **c,** Schematic and experimental timeline for PAGER_TF_ assay. **d,** Summary of candidate antagonist peptides screened in PAGER_TF_, using luciferase gene expression as readout. Antagonist peptides that shifted the EC_50_ more than 10-fold relative to no antagonist (blue dots) were selected for further screening. Peptide sequences and full drug response curves in Extended Data Fig. 1 and 2, respectively. Data representative of *n* = 2 independent experiments. **e**, Schematic showing the use of recombinant TEV protease to relieve auto-inhibition of PAGER. **f**, Bar graph showing firefly luciferase reporter activity of various PAGER_TF_ constructs with or without SalB and TEV treatment. Data representative of *n* = 2 independent experiments. **g**, SalB dose response curves for PAGER_TF_ with or without the arodyn (1-6) antagonist. Data representative of *n* = 4 independent experiments. **h**, Schematic showing the use of surface-expressed GFP on a sender cell to relieve auto-inhibition of PAGER. **i**, Bar graph showing firefly luciferase reporter activity of various PAGER_TF_ constructs with or without SalB and co-culture with surface GFP-expressing HEK 293T cells. Data representative of *n* = 4 independent experiments. **j**, SalB dose response curves for PAGER_TF_ with or without co-culture with surface GFP-expressing sender cells. Data representative of *n* = 1 independent experiment. **k**, Schematic showing the use of soluble GFP to relieve auto-inhibition of PAGER. **l**, Bar graph showing firefly luciferase reporter activity of various PAGER_TF_ constructs with or without SalB and soluble GFP. Data representative of *n* = 3 independent experiments. **m**, SalB dose response curves for PAGER_TF_ with or without soluble GFP. Data representative of *n* = 1 independent experiment. **n**, AlphaFold2 model of PAGER showing how GFP antigen binding sterically occludes the arodyn (1-6) peptide antagonist. **o**, Zoom-in on AlphaFold2 model showing proximity between nanobody antigen binding loops (CDRs) and antagonist peptide fusion site (at N-terminus of nanobody) that leads to steric occlusion of the antagonist peptide (red) by the bound antigen (right, green). **p**, Left, PAGER_TF_ variants with flexible linkers can still be activated by surface GFP. Right, data from experiment performed in HEK 293T cells as in (c). Legend in (q). **q**, Left, PAGER_TF_ variants with flexible GS linkers cannot be activated by soluble GFP due to loss of steric clash. Right, data from experiment performed as in (c). Data in p-q representative of *n* = 1 independent experiment.

In considering the optimal GPCR for the PAGER scaffold, we identified two requirements: 1) that it be activatable by a bioorthogonal small-molecule agonist, and 2) that it be sensitive to the activity of a genetically encodable antagonist that can be fused to the GPCR. We focused on Designer Receptors Exclusively Activated by Designer Drugs (DREADDs^16–18)^, GPCRs that have been engineered to be insensitive to native ligands but be activatable by highly selective drug-like small molecules with minimal activity on endogenous receptors.

We started from the Kappa opioid receptor DREADD (KORD) due to the established activity of peptide antagonists for its parent receptor KOR^18,19^. To read out PAGER activation using a transcriptional reporter, we fused a transcription factor (TF) to KORD via a light-gated protease-sensitive linker, and co-expressed an arrestin-TEV protease (TEVp) fusion (PAGER_TF_; **Extended Data Fig. 1a**)^10,11^. PAGER_TF_ activation triggers arrestin-TEVp recruitment, and if light or furimazine is also present, then the tethered TF will be released by proteolysis, translocate to the nucleus, and drive reporter gene expression (**Fig. 1b–c**). KORD-based PAGER_TF_ was optimized to produce robust expression of a firefly luciferase reporter in response to its small molecule agonist salvinorin B (SalB) (**Extended Data Fig. 1b–d**).

We then screened a library of antagonist peptides to auto-inhibit PAGER_TF_. KOR’s natural agonist, dynorphin, is a 17-amino acid peptide that binds with its N-terminal end buried in the binding pocket of KOR (**Extended Data Fig. 1e**)^20,21^. We selected 24 candidate antagonists based on mutated or truncated variants of dynorphin (sequences in **Extended Data Fig. 1f**) and fused them to the extracellular N-terminal end of KORD, separated by a GFP-specific nanobody (LaG17) and a TEV protease cleavage site (TEVcs) (**Extended Data Fig. 1g**). To ensure cell surface targeting, all constructs were cloned after an IL-2 signal peptide, which leaves no N-terminal scar that could interfere with antagonist function. 16 out of 24 peptides displayed antagonism in the context of PAGER_TF_ by shifting the EC_50_ of the SalB response more than 10-fold (**Fig. 1d**; full dose-response curves in **Extended Data Fig. 2**).

Additionally, for PAGER to function as designed, this peptide antagonism should be reversible. To screen the 16 antagonized constructs for this property, we used extracellular recombinant TEV protease to cleave off the antagonist and relieve antagonism (**Fig. 1e**). We found that all 16 constructs were reversibly antagonized because protease treatment re-sensitized PAGER_TF_ to SalB (**Extended Data Fig. 3**). We selected four constructs that use the dynorphin analogue arodyn^19^ for auto-inhibition, due to their high signal:noise ratio in PAGER_TF_ (**Fig. 1f–g**, **Extended Data Fig. 3**).

With activatable PAGER_TF_ constructs in hand, we next tested for the ability of GFP antigen to relieve auto-inhibition and give robust SalB activation. We performed this test in two separate assays: 1) with surface-expressed GFP introduced via co-culture (**Fig. 1h–j, Extended Data Fig. 4a**), and 2) with soluble recombinant GFP (**Fig. 1k–m, Extended Data Fig. 4b**). All α-GFP PAGER_TF_ constructs responded in varying degrees to both surface-expressed and soluble GFP. We observed that longer arodyn peptides were better antagonists but were also more difficult to remove with GFP antigen, likely due to higher affinity for the receptor. For this reason, we chose the shorter arodyn (1-6) peptide, which displayed sufficient antagonism and the greatest response to GFP, for all PAGER_TF_ constructs going forward.

PAGER was designed so that antigen binding sterically occludes the peptide antagonist, preventing it from occupying the orthosteric site; un-inhibited receptor can then be activated by drug (**Fig. 1n**). We hypothesized that this mechanism is enabled by the proximity between the nanobody’s antigen-binding loops (CDRs 1–3) and N-terminal end, where the peptide antagonist is fused (**Fig. 1o**). This steric activation mechanism should operate for both cell surface and soluble antigens. With cell surface antigens, however, tensile force between PAGER and the target antigen due to cell-cell contact and endocytosis could also displace the auto-inhibitory domain. To probe PAGER’s mechanism, we varied the linker between the nanobody N-terminus and the fused peptide antagonist by inserting one or two copies of a flexible GGGS linker. We hypothesized that inclusion of flexible linkers might relieve any steric occlusion of the antagonist by bound antigen and render PAGER insensitive to soluble antigens. Surface antigens, however, should still be able to activate PAGER using force-based activation. Indeed, when including short 4 or 8 amino acid GS linkers between the nanobody and the peptide antagonist, α-GFP PAGER_TF_ could still be activated by surface GFP antigen (**Fig. 1p**), but its response to soluble GFP antigen was largely abrogated (**Fig. 1q**).

### PAGER can sense and respond to diverse antigens

For PAGER to have programmable antigen-specificity, it should be highly modular. Simply swapping the antigen-binding nanobody in PAGER for another one against a different antigen should produce a new functional receptor capable of sensing and responding to the new antigen of interest. To test this, we first swapped the nanobody in α-GFP PAGER_TF_ for 10 other α-GFP nanobodies and 6 α-mCherry nanobodies. These nanobodies bind to diverse epitopes on the surface of GFP and mCherry and should therefore produce different spatial relationships between bound antigen and fused antagonist peptide. The constructs were first screened for expression, auto-inhibition, and relief of auto-inhibition by TEV protease treatment. All but 2 α-GFP PAGER_TF_ constructs passed this initial screen, as shown in dose-response curves in **Extended Data Fig. 5**. In a subsequent screen with soluble antigen, all 9 remaining α-GFP PAGER_TF_ and all 6 α-mCherry PAGER_TF_ constructs showed clear activation in response to soluble recombinant GFP and mCherry, respectively (**Fig. 2a**; dose response curves in **Extended Data Fig. 6**). We performed a titration with our best α-GFP PAGER_TF_ (LaG2) and our best α-mCherry PAGER_TF_ (LaM8) (**Fig. 2b**). α-GFP PAGER_TF_ could detect soluble recombinant GFP down to a concentration of ∼5 nM, in good agreement with the published K_d_ of the LaG2 nanobody (16–19 nM)^22^, while α-mCherry PAGER_TF_ could detect 100 nM mCherry protein. Neither PAGER was responsive to non-cognate antigen, illustrating the high specificity of these synthetic receptors.

**Figure 2.**
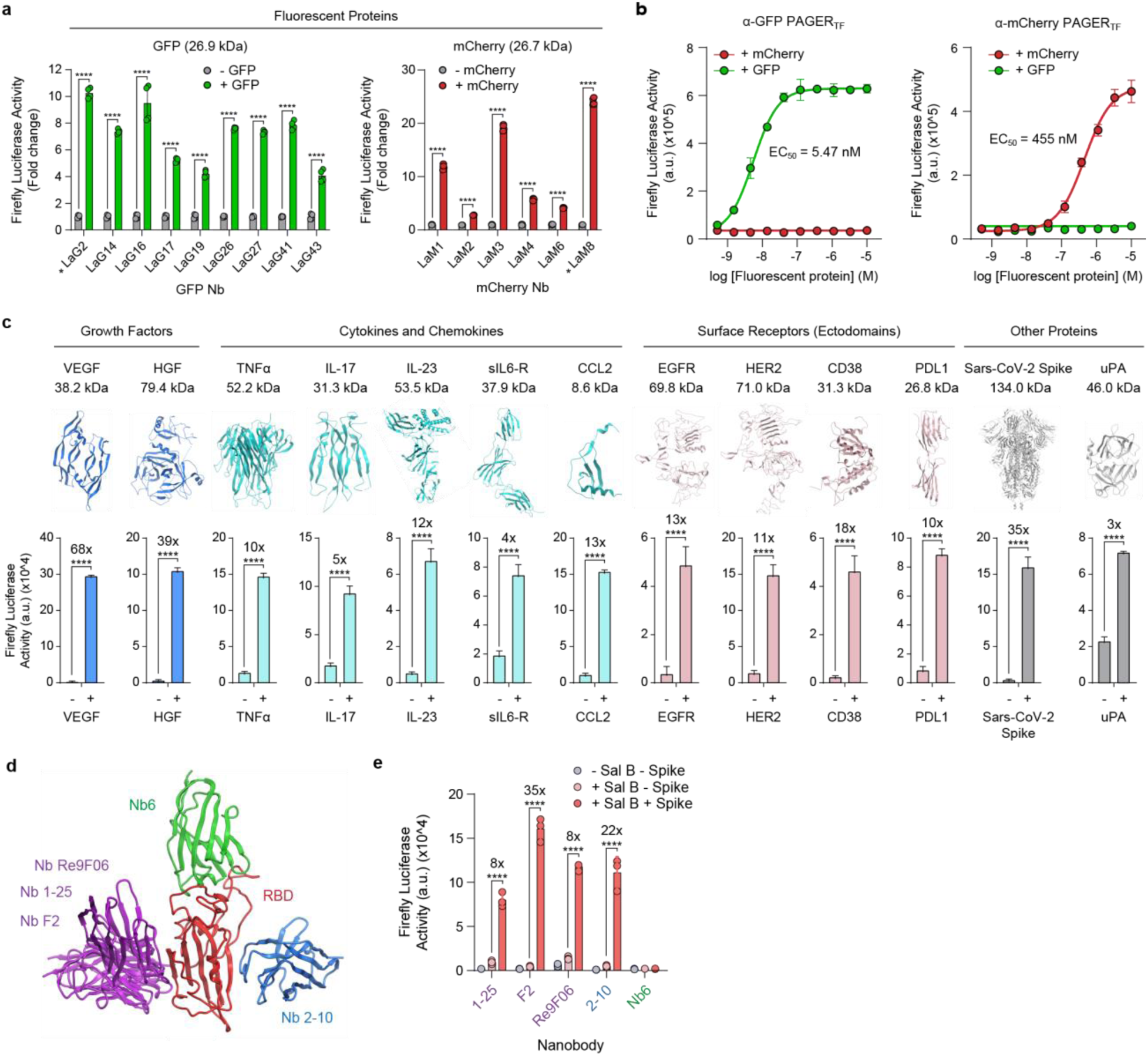
PAGER_TF_ can detect a wide variety of antigens. **a,** Many different GFP and mCherry-binding nanobodies work in PAGER. HEK cells expressing PAGER_TF_ with the indicated nanobodies were stimulated with Salvinorin B and 1 µM soluble GFP or 10µM soluble mCherry, for 15 minutes. Luciferase reporter expression was measured 8 hours later. Data representative of *n* = 2 independent experiments. **b,** mCherry and GFP PAGERs are orthogonal. Data obtained using starred constructs in (a). Add asterisks to LaG2 and Lam8 in (a). Data representative of *n* = 1 experiment. **c,** PAGER_TF_ detects a wide variety of antigens. Only the nanobody was swapped and the rest of the PAGER_TF_ scaffold was not modified. HEK cells were treated with the indicated antigen and SalB for 15 minutes. Antigen and SalB concentrations were determined from initial TEV screen (**Extended Data Fig. 7**) and are reported in Methods. Data representative of *n* = 1-2 independent experiments. **d,** Five nanobodies that bind to the receptor binding domain (RBD) of SARS-CoV2 spike protein as shown^23–26^ were tested in PAGER_TF_. **e,** HEK cells expressing the indicated PAGER_TF_ were treated with 200 nM spike protein and 500 nM SalB for 15 minutes before luciferase measurement 8 hours later. Data representative of *n* = 1 independent experiment.

We then attempted to generate PAGER_TF_’s against many different antigens of various types, sizes, and folds, including growth factors, cytokines, chemokines, receptor tyrosine kinases, other cancer-expressed surface receptors, a viral protein, and a protease, again by simply swapping out the nanobody in PAGER_TF_ for published nanobodies against each new antigen of interest. In this way, we successfully made PAGER_TF_ against GFP, mCherry, VEGF, HGF, TNFα, IL-17, IL-23, sIL-6R, CCL2, EGFR, HER2, CD38, PD-L1, Sars-CoV-2 spike protein, and uPA (**Fig. 2c**); each PAGER was responsive to its cognate antigen.

For Sars-CoV-2 spike protein, many nanobodies have been engineered against its receptor binding domain (RBD), due to the importance of the RBD-ACE2 interaction for viral entry into cells^23–26^. We tested several of these nanobodies in PAGER_TF_ and found five that bind to an epitope on RBD that is accessible in intact trimeric spike protein, and give good cell surface expression and relief of antagonism upon treatment with extracellular TEV protease (**Fig. 2d**). Upon addition of recombinant spike protein to HEK 293T cells, 4 out of 5 PAGER_TF_ constructs produced luciferase reporter gene expression if SalB was also present (**Fig. 2e**). These demonstrations help to illustrate the versatility and broad antigen scope of PAGER.

### Antigen-dependent G protein activation via PAGER

In PAGER_TF_, antigen recognition drives transgene expression as the output. For other applications, it may be desirable for antigen recognition to couple to and modulate endogenous signaling pathways, on a rapid timescale, to control cellular behavior. Because PAGER is based on GPCRs, we wondered whether our platform could convert antigen recognition into rapid activation of endogenous G protein pathways and drive diverse downstream effects on cell behavior that they direct (PAGER_G_; **Fig. 3a**).

**Figure 3.**
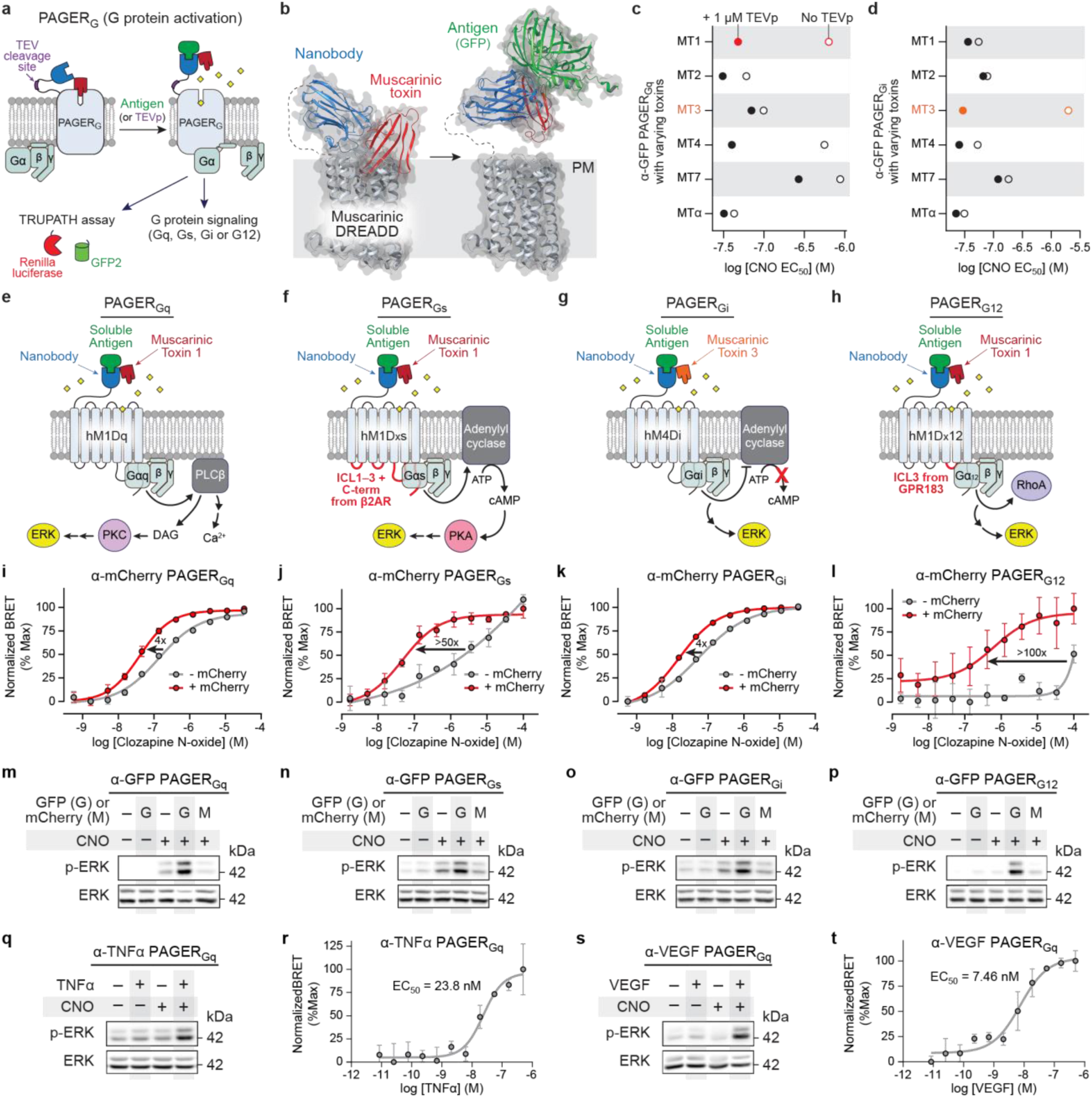
PAGER_G_ couples antigen recognition to activation of endogenous G proteins. **a,** Schematic of PAGER_G_ overview. PAGER_G_ can respond to extracellular antigen, or alternatively TEV protease, similarly to PAGER_TF_ and leads to the activation of selective G protein signaling (Gq, Gs, Gi or G12), which can be measured by TRUPATH BRET assay (see **Extended Data Fig. 7a** for schematic of TRUPATH assay). **b,** AlphaFold2-predicted structure of PAGER_G_ showing auto-inhibition by muscarinic toxin (MT1; red) and release of auto-inhibition upon nanobody (blue) binding to antigen (GFP, green). **c,** Screening toxin variants for auto-inhibition of Gq-coupled PAGER. HEK cells expressing the indicated variant were stimulated with varying concentrations of CNO, with (filled) or without (open) prior TEVp treatment to cleave off the toxin. PAGER_Gq_ activation was measured with TRUPATH BRET assay. Full drug response curves in **Extended Data Fig. 6**. MT1 toxin was best for gating PAGER_Gq_. Data representative of *n* = 1 independent experiment. **d,** Same assay as in (c) but for Gi-coupled PAGER. Full drug response curves in **Extended Data Fig. 6**. MT3 toxin was best for gating PAGER_Gi_. **e–h,** Schematic illustrating the coupling of PAGER_G_ to endogenous intracellular effectors. **i–l,** TRUPATH BRET measurements showing that each α-mCherry PAGER_G_ activates its corresponding G protein in the presence of matched antigen (1 uM mCherry; 1 uM GFP used in negative control). EC50 of each curve is labeled in the plot. Data representative of *n* = 4 independent experiments. **m–p,** Western blots showing phosphorylation of endogenous ERK in response to GFP and CNO activation of each PAGER_G_. HEK cells expressing indicated α-GFP PAGER_G_ was stimulated with 100 nM CNO and 1 μM antigen for 5 min, before cell lysis and analysis. Data representative of *n* = 3 independent experiments. **q–t,** PAGER_Gq_ can be programmed to respond to different antigens. The nanobody in α-GFP PAGER_Gq_ was replaced with nanobodies against TNFα (q–r) and VEGF (s–t). q and s, western blots showing phosphorylation of endogenous ERK after 5-minute stimulation with 100 nM CNO and 100 nM antigen. Data representative of *n* = 3 independent experiments. r and t, TRUPATH BRET measurements showing α-TNFα and α-VEGF PAGER_Gq_ activate Gαq in the presence of matched antigen and CNO. Data representative of *n* = 2 independent experiments.

To explore this possibility, we returned to the other DREADDs, which consist of engineered muscarinic acetylcholine receptors (M1–M5) that activate Gαq (M1, M3, M5) or Gαi (M2, M4)^16^. Additionally, chimeric receptors that activate Gαs or Gα12 have been developed from M3 DREADD^27–29^. These DREADDs no longer bind to their endogenous ligand, acetylcholine, but can be activated by the orthogonal drugs clozapine N-oxide (CNO)^16^ and deschloroclozapine (DCZ)^30^. Though muscarinic GPCRs lack known peptide antagonists, muscarinic toxin (MT) proteins from *Dendroaspis* snakes^31^ can antagonize specific muscarinic receptor subtypes^32,33^. We wondered if these MTs could be used for auto-inhibition to develop PAGER_G_’s (**Fig. 3b**).

Starting with the Gq and Gi DREADDs, we fused a GFP-specific nanobody and a panel of MTs to their N-terminal ends. A TEV protease cleavage site was introduced between the nanobody and DREADD to enable testing for reversible auto-inhibition with addition of extracellular TEV protease as before. To measure PAGER-driven Gq or Gi activation, we used the BRET-based TRUPATH assay^34^. We found that MT1 on hM1Dq DREADD produced the greatest fold-change in Gq recruitment +/– TEV protease, indicating strong and reversible inhibition, while MT3 was the best toxin for gating hM4Di DREADD (**Fig. 3c–d**; dose response curves in **Extended Data Fig. 8**). We next tested for GFP-dependent activation, but to avoid overlap between TRUPATH fluorescence signal and GFP fluorescence, we opted for luciferase gene expression as the readout (**Fig. 1b**). Again, the combination of MT1–hM1Dq and MT3–hM4Di yielded the greatest difference in luminescence +/– GFP (**Extended Data Fig. 9a–c**).

We found that none of the MTs we screened auto-inhibited the hM3Dq DREADD (**Extended Data Fig. 8, Extended Data Fig. 9d–g**), preventing us from building PAGER_Gs_ or PAGER_G12_ from the available hM3Dq-based chimeric Gs and G12 DREADDs^28,29^. The high sequence- and structural-homology between M1 and M3, however, allowed us to build similar chimeras using M1 DREADD. We created Gs and G12-coupled PAGERs by grafting the chimeric components used in Gs- and G12-coupled M3 DREADDs into hM1Dq and utilizing the MT1 toxin for auto-inhibition (**Fig 3e–h, Extended Data Fig. 9h**).

With our panel of PAGER_Gq_, PAGER_Gs_, PAGER_Gi_, and PAGER_G12_ constructs, we next tested generalizability to other antigens. When we replaced the α-GFP nanobody with an α-mCherry nanobody, we observed that soluble mCherry and CNO could activate the corresponding G protein, using a TRUPATH BRET assay (**Fig. 3i–l**). We also measured coupling of PAGER_G_ to endogenous G proteins by blotting for ERK phosphorylation, a conserved downstream response to G protein activation. mCherry and CNO treatment resulted in increased endogenous ERK phosphorylation whereas negative controls omitting either stimulant did not (**Fig. 3m–p**). To test more antigens, we replaced the α-mCherry nanobody in PAGER_Gq_ with an α-TNFα nanobody (ozoralizumab; K_d_ 20.2 pM^35^) or α-VEGF scFv (brolucizumab; K_d_ 28.4 pM^36^). The resulting constructs elicited Gq activation in the presence of cognate antigens as measured with phospho-ERK and TRUPATH BRET assays (**Fig. 3q–t**).

TNFα is an important pro-inflammatory cytokine that is released mainly by macrophages during host defense^37^. Our dose titration showed that α-TNFα PAGER_Gq_ could respond to α-TNFα levels as low as 2 nM. VEGF is also a biologically important signal, released by tumor cells, macrophages, and platelets during angiogenesis and inflammation^38^. α-VEGF PAGER_Gq_ could detect 0.2 nM VEGF, less than the amount released during idiopathic myelofibrosis^39^ for example (1–85 nM). Thus, α-VEGF PAGER_Gq_ may have sensitivity suitable for some *in vivo* applications.

Finally, we studied the mechanism of PAGER_G_ activation. For both α-GFP PAGER_Gq_ and α-GFP PAGER_Gi_, reducing the affinity of MT to the receptor resulted in increased binding of GFP to the α-GFP nanobody, indicating that MT binding to the receptor sterically competes with antigen binding to nanobody (**Extended Data Fig. 9i–j**). Moreover, α-GFP nanobodies with higher reported affinity were better overall at competing with MTs (**Extended Data Fig. 9k**), and they elicited better GFP-dependent activation of PAGER_Gq_ (**Extended Data Fig. 9l**). Interestingly, truncation of MT, or extending the linker between MT and nanobody even by a few amino acids resulted in lack of antagonism or loss of antigen-dependent activation, respectively (**Extended Data Fig. 9m**). These observations together suggest that PAGER_G_ operates by a similar steric occlusion mechanism as PAGER_TF_ and highlight the importance of balancing strong antagonism with facile displacement by antigens of interest.

### Customized cell behaviors driven by PAGER

In nature, heterotrimeric G proteins modulate the concentrations of various intracellular second messengers to drive diverse cellular responses, from cell growth and migration to neuronal inhibition (**Fig. 4a**). We explored the ability of PAGER_G_ to produce customized cellular responses to antigens specified by the nanobody component of PAGER.

**Figure 4.**
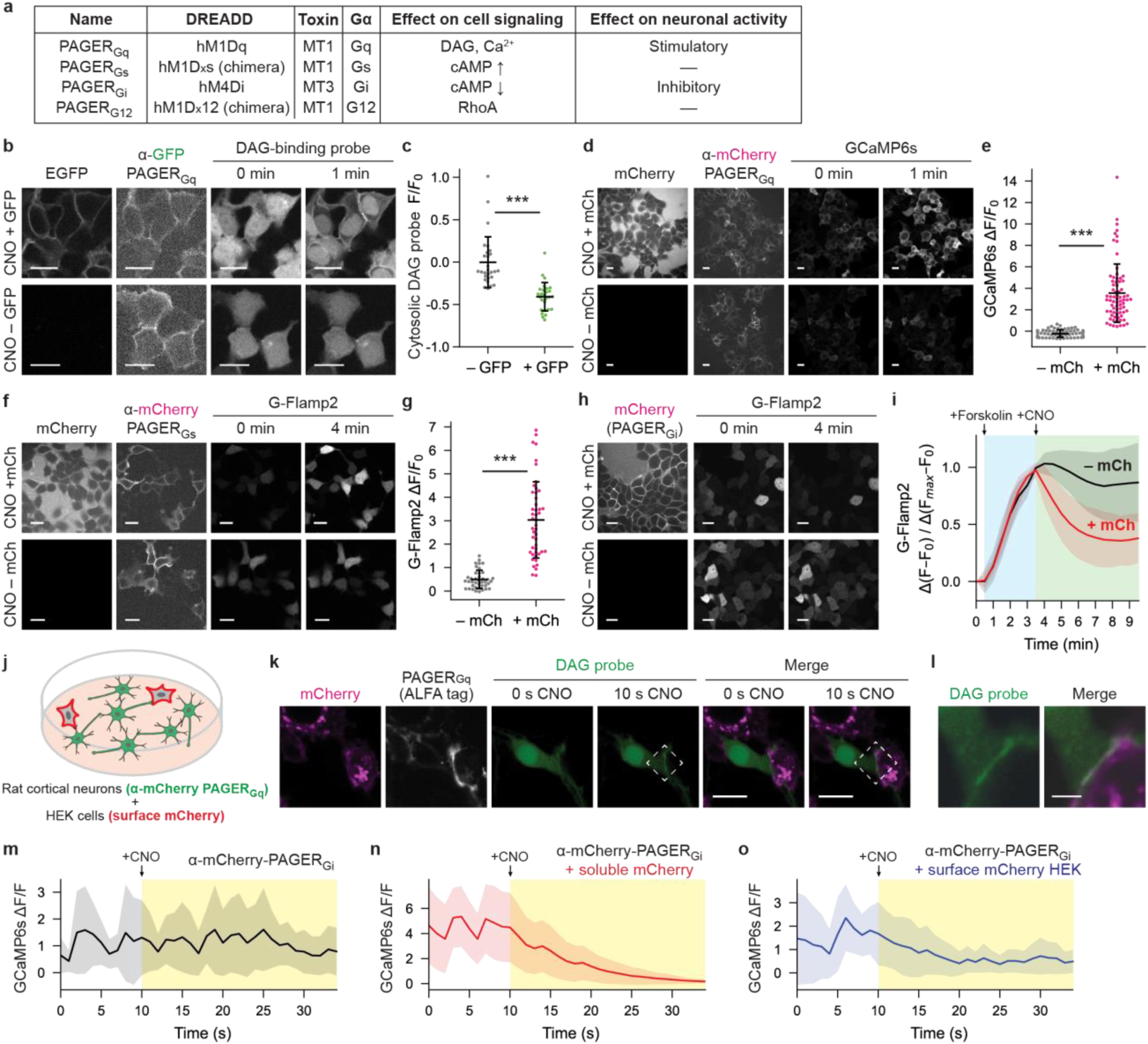
Applications of PAGER_G_ to control cell signaling and behavior. **a,** Table listing the four types of PAGER_G_, the DREADDs and toxins from which they are made, and their effects on downstream signaling. **b,** Antigen-dependent control of endogenous DAG lipid production with PAGER_Gq_. HEK cells expressing α-GFP PAGER_Gq_ and DAG probe (C1_PKCγ_-mCherry^40^) were treated with 1 μM GFP and 100 nM CNO and imaged over time. DAG probe accumulates at the plasma membrane, reflecting Gq-driven phospholipase C activity. **c,** Quantitation of data in (b), comparing depletion of DAG-binding probe from cytosol at the 1-minute timepoint. *n* = 28 and 27 individual cells, respectively, examined over three independent samples. **d,** Antigen-dependent control of cytosolic Ca^2+^ with PAGER_Gq_. HEK cells expressing α-mCherry PAGER_Gq_ and GCaMP6s^41^ were stimulated with 2 μM mCherry and 100 nM CNO and imaged over time. **e,** Quantitation of data in (d), comparing increase in Ca^2+^ sensor (GCaMP6s) intensity at the 1-minute timepoint. *n* = 72 and 73 individual cells, respectively, examined over three independent samples. **f,** Antigen-dependent control of cytosolic cAMP with PAGER_Gs_. HEK cells expressing α-mCherry PAGER_Gs_ and G-Flamp2^42^ reporter were stimulated with 1 μM mCherry and 100 nM CNO and imaged over time. **g,** Quantitation of data in (f), comparing increase in cAMP sensor (G-Flamp2) intensity at the 4-minute timepoint. *n* = 40 and 47 individual cells, respectively, examined over three independent samples. **h,** Antigen-dependent control of cytosolic cAMP with PAGER_Gi_. HEK cells expressing α-mCherry PAGER_Gi_ and G-Flamp2 reporter were stimulated with 1 μM mCherry and 25 nM CNO and imaged over time. **i,** Quantitation of data in (h). *n* = 52 and 42 individual cells, respectively, examined over two independent samples. Averages of Δ(F–F_min_)/(F_max_-F_min_) in G-Flamp2 signal, where F_max_ and F_min_ is the maximum and minimum signals, respectively, recorded in the time window, were plotted, and shaded areas indicate standard deviations. **j,** Schematic for neuron-HEK mixed culture experiment. HEK cells expressing surface mCherry were plated on top of neurons expressing α-mCherry PAGER_Gq_ and either DAG probe (k) or GCaMP6s (m–o). HEK–neuron cultures were stimulated with CNO and imaged over time. **k,** Images from experiment in (j). CNO was added to the final concentration of 100 nM. **l,** Magnification of boxed region in (k) Scale bar, 5 μm. **m–o,** Calcium traces in rat cortical neurons co-expressing α-mCherry PAGER_Gi_ and GCaMP6s, with no antigen (m), with 1 μM mCherry (n), or co-plated with HEK cells expressing surface mCherry (o). CNO was added at t=10 s to the final concentration of 30 nM. Averages of Δ(F–F_base_)/F_base_ in GCaMP6s signal, where F_base_ is the minimum signal recorded in the time window, were plotted, and shaded areas indicate standard deviations (*n* = 12 neurons, data representative of two independent experiments). Scale bar, 20 μm except otherwise noted.

The G protein Gq elevates cytosolic Ca^2+^ and DAG lipid production at the plasma membrane via activation of phospholipase C (PLC). To see if PAGER_Gq_ could activate these endogenous pathways, we transfected HEK cells with α-GFP PAGER_Gq_ and a genetically-encoded DAG probe (mCherry fused to a DAG-binding C1_PKCγ_ domain^40^) that translocates from the cytosol to the plasma membrane upon DAG production. Treatment of cells with GFP and CNO, but not CNO alone, resulted in relocalization of DAG-binding probe from the cytosol to the plasma membrane over 1 minute, consistent with the kinetics of PLC-mediated DAG production^40^ (**Fig. 4b–c**, **Extended Data Fig. 10a**). Using a genetically-encoded Ca^2+^ indicator, GCaMP6s^41^, we similarly verified that α-mCherry PAGER_Gq_ could produce rises in intracellular Ca^2+^ after treatment of HEK cells with mCherry and CNO (**Fig. 4d–e**, **Extended Data Fig. 10b**).

The stimulatory G protein Gs increases levels of cytosolic cAMP, leading to activation of protein kinase A instead of protein kinase C as with Gq. The inhibitory G protein Gi counteracts this by reducing levels of cytosolic cAMP. We prepared HEK cells expressing the recently-developed cAMP indicator G-Flamp2^42^ and either α-mCherry PAGER_Gs_ or α-mCherry PAGER_Gi_. By imaging G-Flamp2, we could observe mCherry-dependent increases in endogenous cAMP when PAGER_Gs_ was expressed and mCherry-dependent decreases in cAMP when PAGER_Gi_ was expressed (**Fig. 4f–i, Extended Data Fig. 10c**).

In neurons, Gq and Gi signaling can result in activation or silencing of neuronal activity, respectively (**Fig. 4a**). Thus Gq- and Gi-coupled DREADDs are widely used in neuroscience to activate or inhibit neuronal subpopulations in response to orthogonal drugs. PAGER_G_ can introduce an additional layer of specificity beyond DREADDs, by conferring antigen-dependence, whereby neuronal activity is controlled by drug *and* by soluble or cell surface antigen. To demonstrate this, we transduced neurons with α-mCherry PAGER_Gq_ and DAG-binding probe. Stimulation of neurons with recombinant mCherry and CNO, but not CNO alone, produced DAG synthesis at the plasma membrane (**Extended Data Fig. 10d–e**). We also prepared a co-culture of α-mCherry PAGER_Gq_-expressing neurons with surface mCherry-expressing HEK cells. Upon addition of CNO, the DAG probe accumulated in regions of the plasma membrane that were in contact with mCherry-expressing HEK, highlighting the potential of PAGER_G_ to confer spatial control of neuronal activity (**Fig. 4k–l, Extended Data Fig. 10f**).

To test PAGER_Gi_ for antigen-dependent control of neuronal inhibition, we transduced neurons with α-mCherry PAGER_Gi_ and the calcium indicator GCaMP6s. Untreated neurons and CNO-only treated neurons exhibited basal calcium spiking activity (**Fig. 4m, Extended Data Fig. 10g**). However, addition of soluble mCherry along with CNO strongly suppressed calcium activity, as did co-culturing with HEK cells presenting surface mCherry (**Fig. 4n–o**, **Extended Data Fig. 10h–i**). These examples show that PAGER_G_ can be used for spatially specific control of neuronal activity with genetically targetable antigens.

### Real-time fluorescent sensors based on PAGER

As a final readout, we explored the use of PAGER for real-time fluorescence detection of antigen binding to cells. GRABs are a class of sensors, widely used in neuroscience, for real time fluorescence detection of neurotransmitters, neuromodulators, and neuropeptides^43–45^. GRABs are designed from GPCRs and install a conformation-sensitive circularly permuted (cp) fluorescent protein between transmembrane segments 5 and 6, to respond to binding of the receptors to their cognate ligands. Many antigens of interest, however, do not have natural GPCRs that can be exploited for development of GRAB-type sensors. In these cases, we wondered whether PAGER, fused to conformation-sensitive cpEGFP, could be used for real-time detection of diverse antigens, producing PAGER_FL_.

First, we attempted to develop a GRAB-type sensor from M4 DREADD (**Extended Data Fig. 11a–b**), from which PAGER_Gi_ was made. Starting from wild-type human M4, we inserted cpEGFP between transmembrane segments 5 and 6, screened many linkers on either side of cpEGFP, optimized the cpEGFP sequence, and obtained an acetylcholine (ACh) sensor, hM4-1.0, with good membrane trafficking, a maximal response of ∼2-fold, and an apparent affinity of ∼225 nM (**Extended Data Fig. 11c–e**). We then introduced the DREADD binding pocket mutations into hM4-1.0 (**Extended Data Fig. 11f**), producing DCZ1.0, which gave 1.7-fold fluorescence turn-on in response to the DREADD ligand DCZ and other designed drugs (**Extended Data Fig. 11g–j**). We confirmed that this sensor does not couple with downstream Gi protein, unlike hM4Di (**Extended Data Fig. 11k**), thus minimizing its potential to perturb native biology.

From DCZ1.0, we produced our first PAGER_FL_ responsive to mCherry antigen (α-mCherry PAGER_FL_) by appending the mCherry-sensitive nanobody LaM6 fused to the inhibitory toxin MT3 (**Fig.5a–b**). Fluorescence measurements showed 300-fold sensitization to DCZ in the presence of mCherry but not non-cognate antigen (BFP) (**Fig. 5c–e**). α-EBFP PAGER_FL_, generated by replacing LaM6 with the GFP- and BFP-binding LaG2 nanobody, showed 130-fold sensitization to DCZ in the presence of EBFP but not mCherry (**Fig. 5f–h**). We then performed time-lapse imaging in HEK cells with sequential addition of DCZ and antigen (**Fig. 5i-k**). While binding of mCherry to α-mCherry PAGER_FL_ had a time constant of t_50_ = 24 s, PAGER_FL_’s response was slower, with a time constant of t_50_ = 3.1 min (**Fig. l**). This lag time reflects PAGER_FL_’s mechanism, which should produce the EGFP-enhancing conformational change only after toxin unbinding/mCherry binding followed by DCZ-mediated activation. Despite the non-instantaneous kinetics, PAGER_FL_ gives much faster readout compared to PAGER_TF_ (translational readout occurs 6–24 h after antigen exposure) and should enable applications not accessible by the latter.

**Figure 5.**
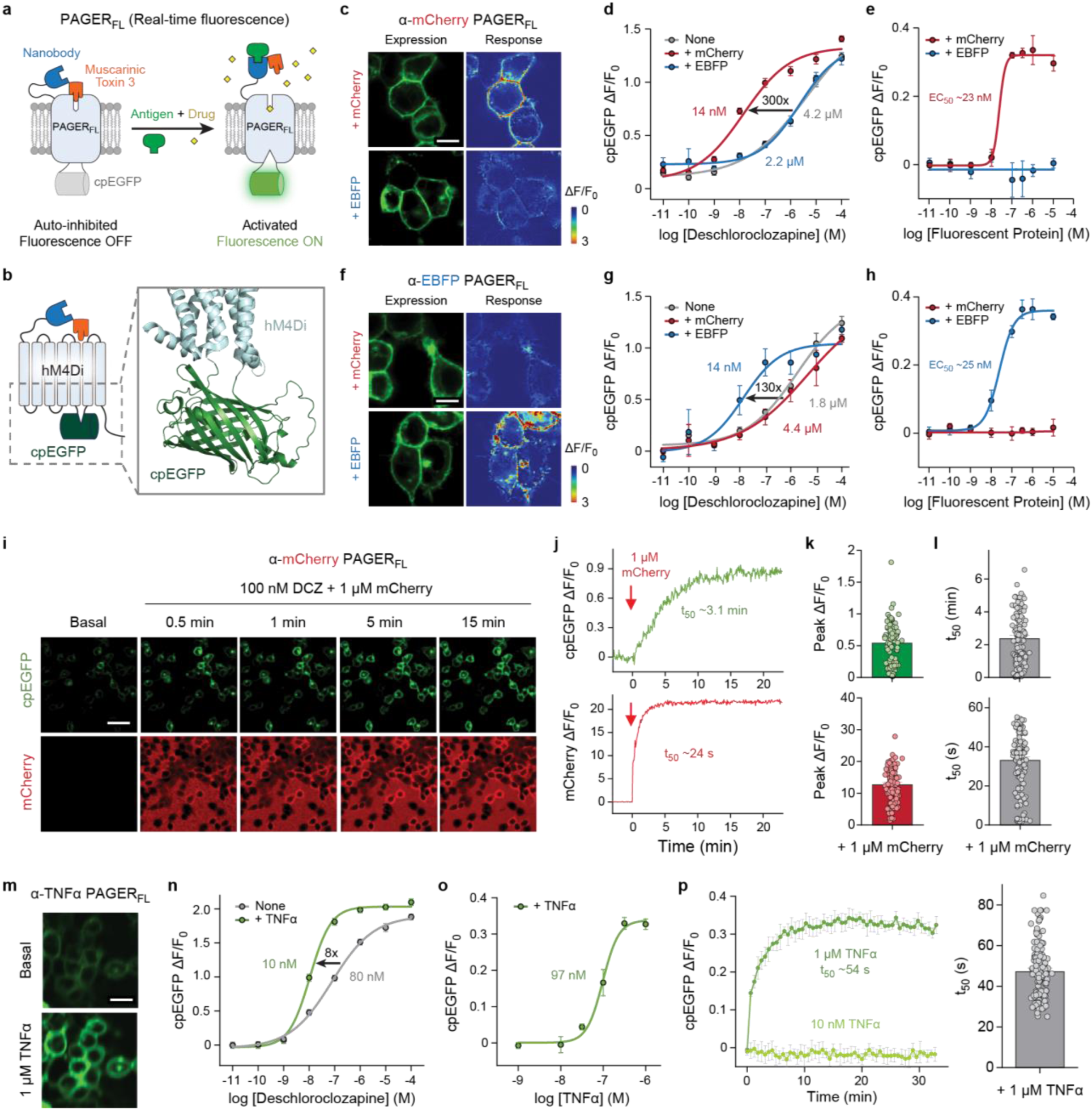
PAGER_FL_ for real-time fluorescence detection of extracellular antigens. **a**, Schematic showing design of PAGER_FL_, based on GRABs^1,2^. A circularly-permuted GFP (cpEGFP) is inserted between transmembrane helices 5 and 6 of PAGER_Gi_. Binding of antigen (green) and deschloroclozapine (DCZ) to PAGER_FL_ triggers a conformational change that increases the fluorescence of fused cpEGFP. **b**, Diagram showing domains of PAGER_FL_, with the modeled structure around the fused cpEGFP shown in zoom. **c**, Representative images of expression and response of α-mCherry PAGER_FL_ to 100 nM DCZ and 1 μM mCherry (antigen) or EBFP (control). Images were collected 30 minutes after addition. Scale bars, 10 μm. **d**, Response of α-mCherry PAGER_FL_ to various concentrations of DCZ, in the presence of 1 μM mCherry (antigen) or EBFP (control). F_0_ is the intensity of sensors in the basal (no DCZ) condition. n = 3 wells containing 100-300 cells per well. **e**, Response of α-mCherry PAGER_FL_ to various concentrations of matched (mCherry) versus mismatched (EBFP) antigen. DCZ was present at 100 nM. F_0_ is the intensity of sensors with 100 nM DCZ addition. n = 3 wells containing 100-300 cells per well. **f-h** are the same as **c-e** but with αEBFP-PAGER_FL_. Scale bars, 10 μm. **i**, Representative confocal images of α-mCherry PAGER_FL_ after addition of 1 μM mCherry in the presence of 100 nM DCZ. Scale bars, 50 μm. **j**, Fluorescence time traces (left) from **i** showing the rate of mCherry and EGFP fluorescence onset after mCherry and DCZ addition. **k**, Summary group data (right) shows the peak response of mCherry and EGFP fluorescence. n = 96 cells from 3 coverslips. **l**, Summary data shows the t_50_ of EGFP fluorescence. n = 144 cells from 4 coverslips. **m**, Representative images of α-TNFα PAGER_FL_, in the basal condition and after addition of 100 nM DCZ and 1 μM TNFα addition. Scale bars, 20 μm. **n**, Response of α-TNFα PAGER_FL_ to various concentrations of DCZ, in the presence of 300 nM TNFα. F_0_ is the intensity of sensors in the basal (no DCZ) condition. n = 3 wells containing 100-300 cells per well. **o**, Response of α-TNFα PAGER_FL_ to various concentrations of TNFα. DCZ was present at 100 nM. F_0_ is the intensity of sensors with 100 nM DCZ addition. n = 3 wells containing 100-300 cells per well. **p**, Fluorescence time traces (left) showing the rate of EGFP fluorescence onset after TNFα addition. DCZ was present at 100 nM. Summary data (right) shows the t_50_ of EGFP fluorescence. n = 137 cells from 3 wells.

To test the ability of PAGER_FL_ to detect other antigens, we prepared a construct using the same TNFα nanobody (Ozoralizumab) that worked well for PAGER_TF_ and PAGER_G_. α-TNFα PAGER_FL_ localized well on the cell membrane when expressed in HEK 293T cells, and the application of 1 μM TNFα elicited fluorescence increase in the presence of DCZ (**Fig. 5m-n**). The response (ΔF/F_0_) and kinetics of α-TNFα PAGER_FL_ were similar to those of α-mCherry PAGER_FL_ (**Fig 5o-p**). Collectively, our results demonstrate that our modular PAGER design can be successfully merged with the GRAB sensor scaffold to enable real-time detection of antigens.

## Discussion

PAGER is a versatile platform for detection of diverse extracellular signals and conversion into a range of user-specified intracellular responses. The high modularity of PAGER, its ability to respond to both surface and soluble antigens, the built-in drug gating, and the ability to dive antigen-dependent transgene expression, G protein activation, or real-time fluorescence, distinguish this platform from other technologies and suggest broad applicability in cell biology and neuroscience.

Early strategies to develop synthetic receptors focused on mutation of natural receptors via site-directed mutagenesis^8,18,46^ or directed evolution^9,16,47^, to alter ligand-specificity. This approach is generally time, labor, and resource intensive and needs to be repeated for every new synthetic receptor being made. An alternative approach alters ligand specificity by replacing the sensing domains of native receptors with other ligand-binding domains such as single-chain antibody variable fragments (scFvs) or nanobodies. This approach creates programmable receptors that can be more easily engineered due to the modularity of the ligand-binding domain. Existing technologies for programmable synthetic receptors capable of target antigen-induced cell signaling include Chimeric Antigen Receptors (CARs)^2^, synthetic Notch receptor (synnotch)^3^, synthetic intramembrane proteolysis receptors (SNIPRs)^48^, Modular Extracellular Sensor Architecture (MESA) receptors^49^, and generalized extracellular molecule sensor (GEMS) receptors^50^.

Despite their considerable contributions to advancing mammalian cell engineering, particularly with regards to therapeutics, these technologies still have important limitations. One prevailing constraint is the general lack of ability to respond to soluble antigens. A few published examples address this by using two non-overlapping binders against a single antigen to drive receptor dimerization^49,50^. However, rarely are suitable pairs of antigen recognition domains available, limiting the generality of this approach. By contrast, PAGER requires a single antigen binding domain, and converts it into a synthetic receptor for detection of soluble *or* tethered antigens, in a single cloning step. PAGER is also differentiated from other technologies by its built-in drug control (which provides temporal specificity and increases signal:background ratio) and the diverse array of outputs that it offers, from transgene expression to fluorescence to control of endogenous G protein pathways.

While many of the antigen binders we tested in PAGER worked on the first try, some failed, and we speculate that the reasons could be: failure to efficiently target to the cell surface, failure to fully auto-inhibit the receptor in the basal state; failure of antigen binding to displace antagonist, poor functionality of the (nanobody) binder when expressed in mammalian cells, or low affinity of binder for antigen. The first two problems can be discovered by screening new PAGERs with extracellular TEV protease to cleave off the antagonist, a recommended first step for future users of this technology. Of the nanobodies that we screened in this study, 83% of the ones that passed the TEV protease test went on to show clear antigen-dependent gating of PAGER_TF_ activity. This suggests that the PAGER backbone is robust and modular, so long as the inserted nanobody does not impair cell surface trafficking or intramolecular binding of the antagonist peptide.

Because of PAGER’s mechanism, we do not expect especially small soluble antigens to work, as these are unlikely to provide the steric interference necessary to relieve antagonism. Smaller antigens may work if their mode of binding interferes with the fused antagonist. The smallest antigen we tested was CCL2 with a size of 8.6 KDa for the monomeric form and 17.2 kDa for the dimer. Oligomerization of antigens increases their functional size and can help promote steric interference and PAGER activation. Almost all our demonstrations used nanobody binders, which have the required architecture for molecular switching between antigen binding and antagonist inhibition. We were also successful with an scFv against VEGF (Fig. 3s–t), and we speculate that monobodies and V_H_ domains could also work in PAGER due to their similar folds to nanobodies, which bring the antigen binding loops close to their N-terminal ends.

Though PAGER_TF_, PAGER_G_, and PAGER_FL_ are highly related, we noticed important differences between PAGER_TF_, which is based on the kappa opioid receptor and uses the arodyn peptide for auto-inhibition, and the latter two, which are based on the muscarinic receptors and use protein toxins for auto-inhibition. First, PAGER_G_ tends to require higher-affinity antigen binders to relieve antagonism, probably because muscarinic toxins bind with higher affinity than truncated arodyn peptide. Second, PAGER_TF_ works best with nanobody binders, whereas PAGER_G_ seems to accept both nanobodies and scFv binders, which are twice the size, consisting of V_H_ and V_L_ domains joined by a flexible linker. Two explanations are plausible. The crystal structure of the kappa opioid receptor in complex with dynorphin^21^ (PDB: 8F7W) shows that the first 6 amino acids are deeply buried within the ligand binding pocket. Thus, a fused nanobody binder would be pushed closely against the receptor. The larger size of scFv may not accommodate this. The second possible explanation relates to the hydrophobic nature of the arodyn antagonist peptide (FFFRLR). A larger scFv binder, with hydrophobicity at the interface of V_H_ and V_L_, provides more opportunities for the arodyn peptide to engage in non-specific interactions, instead of binding to the orthosteric site and properly inhibiting the GPCR.

In future work, the PAGER platform could be improved and extended in a number of ways. Instead of requiring 3 separate transgenes, the design of PAGER_TF_ could be simplified to require only 2, similar to how we simplified our calcium integrator FLARE^51^. For *in vivo* applications, it would be beneficial to remove the requirement for light or furimazine^11^ to uncage PAGER_TF_’s LOV domain. To ensure full orthogonality *in vivo*, we could also mutate the intracellular loops of PAGER_TF_ to abolish recognition of G_i_, drawing inspiration from a DREADD that recruits arrestin without coupling to any of the G proteins^52^. Finally, we note that PAGER’s use of antigen binding to relieve auto-inhibition could be extended to other classes of proteins, for example to produce antigen-gated enzymes or ion channels.

## Materials and methods

### Plasmid constructs and cloning

Constructs used for transient expression in HEK293T cells were cloned into the pAAV viral vector. For stable expression, the constructs were cloned into the pCDH viral vector. For all constructs, standard cloning procedures were used. PCR fragments were amplified using Q5 polymerase (NEB). Vectors were digested with NEB restriction enzymes and ligated to gel-purified PCR products using T4 ligation, Gibson, NEB HiFi, or Golden Gate assembly. Ligated plasmids were introduced into competent XL1-Blue, NEB5-alpha, or NEB Stable bacteria via heat shock transformation.

### HEK293T cell culture

HEK293T cells were obtained from ATCC (tested negative for mycoplasma) and cultured as monolayers in complete growth media: Dulbecco’s Modified Eagle Medium (DMEM, Corning) containing 4.5 g/L glucose and supplemented with 10% Fetal Bovine Serum (FBS, VWR), 1% (v/v) GlutaMAX (Gibco), and 1% (v/v) Penicillin-Streptomycin (Corning, 5000 units/mL of penicillin and 5000 μg/mL streptomycin), at 37°C under 5% CO_2_. For experimental assays, cells were grown in 6-well, 12-well, 24-well, or 96-well plates pretreated with 20 µg/mL human fibronectin (Millipore) for at least 10 min at 37°C.

### HEK293T cell transient transfection

A 1 mg/mL solution of PEI Max (Polysciences, catalog no. 24765) was prepared for transient transfection as follows. Polyethylenimine (PEI, 500 mg) was added to 450 mL of Milli-Q H_2_0 in a 500 mL glass beaker while stirring with a stir bar. Concentrated HCL was added dropwise to the solution until the pH was less than 2.0. The PEI solution was stirred until PEI was dissolved (∼2-3 hours). Concentrated NaOH was then added dropwise to the solution until the pH was 7.0. The volume of the solution was then adjusted to 500mL, filter-sterilized through a 0.22-μm membrane, and frozen in aliquots at -20°C. Working stocks were kept at 4°C for no more than 1 month.

For transient transfection, HEK293T cells were grown in 6-well, 12-well, or 24-well plates pretreated with 20 µg/mL human fibronectin (Millipore) for at least 10 min at 37°C. Cells were grown to a confluency of ∼70-90% prior to transfection. DNA transfection complexes were made by mixing DNA and 1 mg/mL PEI solution in serum-free DMEM at a 1 μg DNA: 5 μL PEI (1 mg/mL): 100 μL serum-free DMEM. Complexes were allowed to form for 20 min at room temperature. After 20 min, complexes were diluted in complete DMEM up to the growth volume per well size (2.5 mL for 6-well, 1 mL for 12-well, and 500 μL for 24-well). The entire well volume of the HEK293T cells was replaced with the diluted complexes and allowed to transfect cells at 37°C for 5-24 hours. Complete transfection protocols including amounts of DNA and length of transfection are described for each experiment below.

### Firefly Luciferase Reporter PAGER-SPARK experiments

HEK293T cells were plated in human fibronectin-coated 6-well dishes at a density of 750,000 cells per well and allowed to grow overnight (∼18 hours) at 37°C until they reached ∼70-90% confluency. After ∼18 hours, the cells were transfected with 350 ng of the indicated Antagonist-Nanobody-GPCR-eLOV-TEVcs-Gal4 (PAGER_TF_) receptor plasmid, 100 ng of NanoLuc-βarrestin2-TEVp plasmid, and 150 ng of UAS-Firefly Luciferase (FLuc) plasmid. Cells were transfected for 5 hours at 37°C. After 5 hours of transfection, cells from each well were lifted and resuspended in 6 mL of complete DMEM to make an ∼400,000 cells per mL single cell suspension, and 100 μL of cell suspension (∼40,000 cells) was plated per well in a human fibronectin-coated white, clear bottom 96-well plate in triplicate. Plates were wrapped in aluminum foil to protect them from light and incubated at 37°C overnight (∼18 hours). After ∼18 hours, cells should be stimulated.

Stimulation should take place in a dark room under red light (red light does not open LOV domain). Stimulation solutions were optimized for each given antigen and PAGER receptor. Unless otherwise indicated, PAGERs were stimulated as follows: GFP(LaG17/LaG2/LaG16)-PAGERs were stimulated with 1 μM GFP, 1 μM salvinorin B, and 1x furimazine; mCherry(LaM6)-PAGERs were stimulated with 1 μM mCherry, 1 μM salvinorin B, and 1x furimazine; VEGF(Nb35)-PAGERs were stimulated with 500 nM VEGF, 500 nM salvinorin B, and 1x furimazine; HGF(Nb1E2)-PAGERs were stimulated with 250 nM HGF, 250 nM salvinorin B, and 1x furimazine; TNFα(Ozoralizumab)-PAGERs were stimulated with 500 nM TNFα, 250 nM or 500 nM salvinorin B, and 1x furimazine; IL17(Sonelokimab)-PAGER was stimulated with 500nM IL-17, 100 nM salvinorin B, and 1x furimazine, IL23(Nb22E11)-PAGER was stimulated with 250 nM IL-23, 500 nM salvinorin B, and 1x furimazine; sL6R(Voberilizumab)-PAGER was stimulated with 500 nM sIL-6R, 500 nM salvinorin B, and 1x furimazine; CCL2(Nb8E10)-PAGER was stimulated with 1 μM CCL2, 500 nM salvinorin B, and 1x furimazine; EGFR(NbEgB4)-PAGERs were stimulated with 500 nM EGFR ECD, 100nM or 250 nM or 500 nM salvinorin B, and 1x furimazine; HER2(Nb2Rs15d)-PAGERs were stimulated with 500 nM HER2 ECD, 500 nM or 1 μM salvinorin B, and 1x furimazine; CD38(NbMU375)-PAGER was stimulated with 1 μM CD38 ECD, 500 nM salvinorin B, and 1x furimazine; PDL1(KN035)-PAGER was stimulated with 1 μM PD-L1 ECD, 500 nM salvinorin B, and 1x furimazine; SarsCoV2-RBD(NbF2)-PAGER was stimulated with 200 nM Sars-CoV-2 Spike protein, 500 nM or 1 μM salvinorin B, and 1x furimazine; uPA(Nb4)-PAGER was stimulated with 500 nM uPA, 250 nM or 500 nM salvinorin B, and 1x furimazine.

For stimulations, growth media was removed from the 96-well plate by flicking off and dabbing excess on paper towel. To initiate stimulation, 100 μL stimulation solution was added to each well for a total of 15 min. After 15min, stimulation solution was removed by flicking off and dabbing excess on a paper towel, and 100uL of complete DMEM was added back to each well. Plates were again wrapped in aluminum foil and placed in 37°C incubator for 8 hours. After 8 hours post-stimulation, media was removed from 96-well plate by flicking off and dabbing excess on paper towel. Wells were washed once with 125 μL DPBS, and then 50 uL of 1x Bright-Glo (2x diluted 1:1 in DPBS; Promega) was added to each well and incubated for 1 min. After 1 min, firefly luciferase luminescence was measured using a Tecan Infinite M1000 Pro plate reader using the following parameters: 1000 msec acquisition time, green-1 filter (520-570 nm), 25°C linear shaking for 10 sec.

In some experiments where indicated, exogenous ambient room white light was used to uncage the LOV domain instead of furimazine-dependent NanoLuc BRET. In these experiments, furimazine was not included in the stimulation solutions; all else remained the same.

In some experiments where indicated, salvinorin B or antigen dose response curves were analyzed. In these experiments, the concentrations of salvinorin B or antigen were included in the stimulation solutions at different concentrations according to those indicated; all else remained the same.

### HEK293T co-culture for trans assays

For trans assays using HEK293T cells, cells were cultured in 6-well and 12-well plates as described above. Receiver cells in 12-well plates were transfected with 140 ng of the indicated pAAV-Antagonist-Nanobody-GPCR-eLOV-TEVcs-Gal4 receptor plasmid, 40 ng of pAAV-NanoLuc-βarrestin2-TEVp plasmid, and 60 ng of pAAV-UAS-Firefly Luciferase (FLuc) plasmid. Sender cells in 6-well plates were transfected with 2 μg of pAAV-GFP-PDGFR transmembrane domain (surface-expressed GFP). Cells were transfected for 5 hours in a 37°C incubator. After 5 hours, cells were lifted with trypsin, washed with DPBS, and resuspended in 6.25 mL or 2.5 mL of complete DMEM per well for 6-well and 12-well plates, respectively. Sender and receiver cells were mixed at a 4:1 sender:receiver ratio and then 100uL of cell mixtures were plated into 96-well white, clear-bottom microplates at a density of 40,000 cells/well. Plates were wrapped in aluminum foil and incubated for ∼18 hr in a 37°C incubator, then the stimulations and luciferase reporter assay were performed as described above.

### TRUPATH G protein activation BRET assay

HEK293T cells were plated in human fibronectin-coated 6-well dishes at a density of 1,250,000 cells per well and allowed to adhere and grow for 2-4 hours at 37°C. After ∼2-4 hours, the cells were transfected with 250 ng of the indicated G protein-PAGER receptor plasmid, 250 ng of the corresponding Gα-RLuc8 TRUPATH plasmid (Gαi1-RLuc8, GαsS-RLuc8, Gαq-RLuc8, or Gα12-RLuc8), 250 ng of Gβ3 TRUPATH plasmid, and 250 ng Gγ9-GFP2 TRUPATH plasmid. Cells were incubated at 37° and transfection was allowed to proceed for ∼20-24 hours. After transfection, cells from each well were lifted and resuspended in 6 mL of complete DMEM to make an ∼400,000 cells per mL single cell suspension, and 100 μL of cell suspension (∼40,000 cells) was plated per well in a human fibronectin-coated white, clear bottom 96-well plate in triplicate. Plates were incubated at 37°C for ∼20-24 hours. For protease activation of G protein-PAGERs, cells were treated with 1μM TEV protease for 90 min followed by stimulation with various concentrations of CNO and 20μM CTZ400a (substrate for TRUPATH assay) for 5 min before reading out BRET. For antigen activation of G protein-PAGERs, cells were treated with 1μM mCherry for 15 min followed by stimulation with various concentrations of CNO and 20μM CTZ400a for 5 min before reading out BRET. BRET was readout using a Tecan Infinite M1000 Pro plate reader using the following parameters: Filter 1 Magenta (370 to 450 nm), 500 ms integration time; Filter 2 Green (510 to 540 nm), 500 ms integration time; 25°C. Data is presented as NET BRET and displayed as scatter plots with variable slope (four parameter) non-linear regression lines.

### Lentivirus generation and stable cell line generation

To generate lentivirus, HEK293T cells were cultured in T25 flasks and transfected at ∼70% confluency with 2.5 μg of the pCDH lentiviral transfer vector of interest and packaging plasmids psPAX2 (1.25 μg) and pMD2.g (1.25 μg) with 25 µL of polyethyleneimine (PEI, 1 mg/mL; Polysciences). Approximately 72 hours post-transfection, the cell medium was collected and centrifuged for 5 min at 300 x g to remove cell debris. Media containing lentivirus was used immediately for transduction or was aliquoted into 0.5 mL aliquots, flash-frozen in liquid nitrogen, and stored at –80°C for later use. Frozen viral aliquots were thawed at 37°C prior to infection.

HEK293T cells were plated on 6-well human fibronectin-coated plates. When cells reached ∼70-90% confluency, cells were transduced with lentivirus for 1-3 days. The cells were then lifted and replated om a T25 flask, and stably expressing cells were selected for in complete DMEM containing 1μg/mL puromycin for at least 1 week. Cells were split and expanded when they reached ∼80-90% confluency. Cells were maintained under this puromycin selection until the time of experiments. Construct expression was confirmed by flow cytometry, immunofluorescence imaging, or functional characterization.

### Quantification of p-ERK by western blotting

Antigen was added at indicated concentration to HEK 293T cells stably expressing PAGERG. 3 min later, 300 nM of CNO in 500 μL blank DMEM was added to a final concentration of 100 nM, and the cells were incubated for another 3 min. Cells were then lysed with RIPA lysis buffer supplemented with protease and phosphatase inhibitors (50 mM Tris-HCl, pH 7.4, 150 mM NaCl, 1% Triton X, 0.5% sodium deoxycholate, 0.1% SDS, 1 mM EDTA, 1× Halt Protease Inhibitor Cocktail from Thermo Scientific, 1x Phosphatase Inhibitor Cocktail from Cell Signaling Technology). After sonication and centrifugation, the lysate supernatants were mixed with 6× Laemmli sample buffer to prepare the sample for western blotting. The membrane was blotted with 1:1,000 dilutions of antibodies for phospho-p44/42 MAPK (phospho-Erk1/2; Cell Signaling Technology #9101), p44/42 MAPK (Erk1/2; Cell Signaling Technology #9107), and β-Tubulin (Cell Signaling Technology #86298).

### Fluorescence imaging

Confocal imaging was performed on a Zeiss AxioObserver inverted confocal microscope with 10x and 20x air objectives, and 40x and 63x oil-immersion objectives, outfitted with a Yokogawa spinning disk confocal head, a Quad-band notch dichroic mirror (405/488/568/647), and 405 (diode), 491 (DPSS), 561 (DPSS) and 640 nm (diode) lasers (all 50 mW). The following combinations of laser excitation and emission filters were used for various fluorophores: GFP (491 laser excitation; 528/38 emission), mCherry/Alexa Fluor 568 (561 laser excitation; 617/73 emission), Alexa Fluor 647 (647 excitation; 680/30 emission), and differential interference contrast (DIC). Acquisition times ranged from 100 to 500 ms. All images were collected and processed using SlideBook (Intelligent Imaging Innovations).

The Opera Phenix high-content screening system (PerkinElmer) was utilized for GRAB sensors imaging, equipped with a 20x 0.4-NA objective, a 40x 0.6-NA objective, a 40x 1.15-NA water-immersion objective, a 488-nm laser, and a 561-nm laser. GFP and RFP signals were collected using a 525/50 nm emission filter and a 600/30 nm emission filter, respectively. HEK293T cells expressing the GRAB_DCZ1.0_ sensor were imaged before and after adding specified drugs at different concentrations while being bathed in Tyrode’s solution. The change in fluorescence intensity of GRAB_DCZ1.0_ was determined by calculating the change in the GFP/RFP ratio and expressed as ΔF/F_0_.

### Fluorescent DAG assay

An mCherry-based fluorescent DAG biosensor was made by C-terminally tagging mCherry to the C1_PKCγ_ from Addgene plasmid #21205^40^ and cloning into the pCDH lentivirus backbone. HEK 293T cells stably co-expressing α-GFP (LaG16) PAGER_Gq_ and C1_PKCγ_-mCherry were incubated in 1:1000 anti-ALFA–AlexaFluor647 and 1 μM EGFP for 3 min. Cells were then located under the microscope and time-lapse images were obtained every 4 s, and 1 mL of 150 nM CNO was added (to a final concentration of 100 nM) between the first and the second frame. Images at the first time frame (t=0) and 15th time frame (t=60s) were used for analysis. Images were analyzed using ImageJ software. Region of interests (ROIs) were manually added to images and the mean of cytosolic mCherry fluorescence in each cell was quantified to plot the time course of C1_PKCγ_-mCherry signal. The difference in mean fluorescence compared to the initial fluorescence (ΔF/F0) was calculated and used for statistical comparison.

### GCaMP6s calcium assay

A GFP-based fluorescent calcium biosensor was made by cloning GCaMP6s from Addgene plasmid #40753^41^ and cloning into the pCDH lentivirus backbone. HEK 293T cells stably co-expressing α-mCherry (LaM6) PAGER_Gq_ and GCaMP6s were incubated in 1:1000 anti-ALFA– AlexaFluor647 and 1 μM mCherry for 3 min. Cells were then located under the microscope and time-lapse images were obtained every 4 s, and 1 mL of 150 nM CNO was added (to a final concentration of 100 nM) between the first and the second frame. Images at the first time frame (t=0) and 15th time frame (t=60s) were used for analysis. Images were analyzed using ImageJ software. Region of interests (ROIs) were manually added to images and the mean of cytosolic GFP fluorescence in each cell was quantified to plot the time course of GCaMP6s signal. The difference in mean fluorescence compared to the initial fluorescence (ΔF/F0) was calculated and used for statistical comparison.

### G-Flamp2 cAMP assay

A GFP-based fluorescent cAMP biosensor was made by cloning G-Flamp2 from Addgene plasmid # 192782^42^ and cloning into the pCDH lentivirus backbone.

For PAGER_Gs_, HEK 293T cells stably expressing G-Flamp2 were transiently transfected with α-mCherry (LaM6) PAGER_Gs_, and the cells were incubated in 1:1000 anti-ALFA– AlexaFluor647 and 1 μM mCherry for 3 min. Cells were then located under the microscope and time-lapse images were obtained every 10 s, and 1 mL of 60 nM CNO was added (to a final concentration of 50 nM) at t=30s. Images at the CNO addition (t=30s) and the last acquisition (t=240s) were used for analysis. Images were analyzed using ImageJ software. Region of interests (ROIs) were manually added to images and the mean of cytosolic GFP fluorescence in each cell was quantified to plot the time course of G-Flamp2 signal. The difference in mean fluorescence compared to the initial fluorescence (ΔF/F0) was calculated and used for statistical comparison.

For PAGER_Gi_, HEK 293T cells stably expressing G-Flamp2 and α-mCherry (LaM6) PAGER_Gi_ were incubated in 1:1000 anti-ALFA–AlexaFluor647 and 1 μM mCherry for 3 min. Cells were then located under the microscope and time-lapse images were obtained every 20 s, and 500 μL of 2 μM Forskolin was added (to a final concentration of 1 μM) at t=30s, followed by 1 mL of 50 nM CNO (to a final concentration of 25 nM) at t=210s (3.5 min). Images were analyzed using ImageJ software. Region of interests (ROIs) were manually added to images and the mean of cytosolic GFP fluorescence in each cell was quantified to plot the time course of G-Flamp2 signal.

### AAV1/2 generation

To generate supernatant AAV, HEK 293T cells were cultured in 6-well plate and transfected at approximately 80% confluency in opti-MEM reduced serum medium (Gibco). Per each well, the AAV vector containing the gene of interest (360 ng) and AAV packaging/helper plasmids AAV1 (180 ng), AAV2 (180 ng), and DF6 (720 ng) incubated with 10 μL PEI in 200 μL opti-MEM were used for transfection. After 20 h, the cell medium was replaced with complete DMEM. The cell medium containing the AAV was harvested 48 h post transfection and filtered using a 0.45 μm filter.

### Neuronal activity assay

All procedures were approved and carried out in compliance with the Stanford University Administrative Panel on Laboratory Animal Care, and all experiments were performed in accordance with relevant guidelines and regulations. Before dissection, 35 mm glass bottom dishes (CellVis) were coated with 0.001% (w/v) poly-l-ornithine (Sigma-Aldrich) in DPBS (Gibco) at room temperature overnight, washed three times with DPBS, and subsequently coated with 5 μg/ml of mouse laminin (Gibco) in DPBS at 37 °C overnight. Cortical neurons were extracted from embryonic day 18 Sprague Dawley rat embryos (Charles River Laboratories, strain 400) by dissociation in Hank’s balanced salt solution with calcium and magnesium (Gibco). Cortical tissue was digested in papain according to the manufacturer’s protocol (Worthington), then 5 × 10^5^ cells were plated onto each dish in neuronal culture medium at 37 °C under 5% CO2. The neuronal culture medium is neurobasal (Gibco) supplemented with 2% (v/v) B27 supplement (Life Technologies), 0.5% (v/v) fetal bovine serum, 1% (v/v) GlutaMAX, 1% (v/v) penicillin-streptomycin, and 1% (v/v) sodium pyruvate (Gibco, 100 mM). On DIV 3 and DIV 6, Half of the media was removed from each dish and replaced with neuronal culture medium. On DIV 6 after the media change, each well was infected with 35 μL of AAV1/2 (10 μL of GCaMP6s AAV and 25 μL of α-mCherry (LaM6) PAGER_Gi_ AAV). Neurons were wrapped in aluminum foil and allowed to express in the incubator. On DIV 13, cells were preincubated in HBSS for 10 min, and then incubated in 1:1000 anti-ALFA–AlexaFluor647 and 1 μM mCherry in HBSS for 3 min. Cells were then located under the microscope and time-lapse images were obtained every 1 s, and 1 mL of 50 nM CNO was added (to a final concentration of 33 nM) at t=10s. Images were analyzed using ImageJ software. Region of interests (ROIs) were manually added to images and the mean of cytosolic GFP fluorescence in each cell was quantified to plot the time course of GCaMP6s signal.

### The development of GRAB_DCZ_ sensors

We chose human M4R as the sensor scaffold and embarked on a systematic optimization process. This process included screening and optimizing the insertion sites, the amino acid composition of the linker, and the critical residues in cpEGFP to enhance the maximum response and fluorescence of sensors. Subsequently, specific DCZ sensors were developed by introducing binding pocket mutations based on these sensors.

① ICL3 replacement: We replaced the ICL3-cpEGFP of the previously developed GRAB_gACh_ sensor^1^ with the corresponding ICL3 of hM4R. A replacement library was generated using 9 sites (S5.62 to H5.70) from the N-terminus and 5 sites (T6.34 to F6.38) from the C-terminus. After screening, we created a prototype ACh sensor named hM4-0.1, which exhibited a 100% ΔF/F fluorescence response to 100 μM ACh. The replacement sites of hM4-0.1 are located between R5.66 and T6.36 in hH4R.
② Linker optimization: The amino acid composition of the linker was found to be critical to the sensor’s dynamic range. We performed site-saturation mutagenesis on 6 residues of the linker. Through this process, we identified a variant named hM4-0.5, with an R5.66L mutation, which resulted in a ∼130% increase in ΔF/F0.
③ cpEGFP optimization: Building on our screening experience in developing GRAB sensors^2,3^, we selected 4 residues in the cpEGFP for individual randomizations. This led to the development of the hM4-1.0 sensor with an H18I mutation, showing a maximal response of ∼350% to 100 μM ACh.
④ Binding pocket mutations: To develop specific DCZ sensors, we introduced Y3.33C and A5.46G mutations^16,54^ based on the hM4-1.0, resulting in the creation of DCZ1.0, which exhibited a ∼150% response to 1 μM DCZ.

### Mini G protein luciferase complementation assay

HEK293T cells were cultured in 6-well plates until they reached 60–70% confluence. At this point, the specified wild-type receptor or sensor, along with the corresponding LgBit-mGi construct, were co-transfected into the cells. Around 24–36 hours post-transfection, the cells were detached using a cell scraper, suspended in PBS, and then transferred to 96-well plates (white with a clear flat bottom) containing Nano-Glo Luciferase Assay Reagent (Promega) diluted 1000-fold in PBS at room temperature. Following this, solutions with varying DCZ concentrations and 1 μM antigens were added to the wells. After a 10-minute reaction in the dark at room temperature, luminescence was measured using a VICTOR X5 multi-label plate reader (PerkinElmer).

### Quantification and Data Analysis

All graphs were created using GraphPad Prism 9 or matplotlib (Python). Error bars represent standard deviation unless otherwise noted. For comparison between two groups, p-values were determined using two-tailed Student’s t tests. For multiple comparisons, p-values were determined using two-way ANOVA with Tukey’s multiple comparisons test to adjust for multiple comparisons. * p < 0.05; ** p < 0.01; *** p < 0.001; **** p < 0.0001; n.s. - not significant.

## Data availability

All data supporting the findings of this study are available within the paper and its Supplementary Information. No datasets were generated or analyzed during the current study.

## Acknowledgements

We are grateful to the Chan Zuckerberg Biohub - San Francisco, Emerson Collective, Stanford Cancer Institute, Wu Tsai Neurosciences Institute of Stanford University (Neuro-omics grant to A.Y.T.), and NIH/NCI (F32CA257159 to N.A.K.) for support of this work. This work was also supported by the National Natural Science Foundation of China (31925017 to Y.L.), and the New Cornerstone Science Foundation through the New Cornerstone Investigator Program and the XPLORER PRIZE (to Y.L.). R.T. was supported by the Life Sciences Research Foundation Fellowship (sponsored by Astellas Pharma) and JSPS Overseas Research Fellowship. We acknowledge Dr. Sungmoo Lee and Prof. Michael Lin for providing rat cortices. This work is dedicated to Tony Kalogriopoulos, may his memory be eternal.

## Author Contributions

N.A.K. and A.Y.T. conceived this project. N.A.K. and A.Y.T. designed experiments and analyzed data for PAGER_TF_. R.T., N.A.K. and A.Y.T. designed experiments and analyzed data for PAGER_G_. Y.Y. and Y.L. designed experiments and analyzed data for PAGER_FL_. M.A.R. designed the TEV protease assay to screen for reversible inhibition of PAGER. N.A.K. performed all experiments for PAGER_TF_ and TRUPATH assays in PAGER_G_, R.T. performed all other experiments in PAGER_G_, and Y.Y. performed all experiments in PAGER_FL_. N.A.K., R.T. and A.Y.T. wrote the manuscript with input and edits from all authors.

