## Supplementary Information for "Synthetic G protein-coupled receptors for programmable sensing and control of cell behavior"

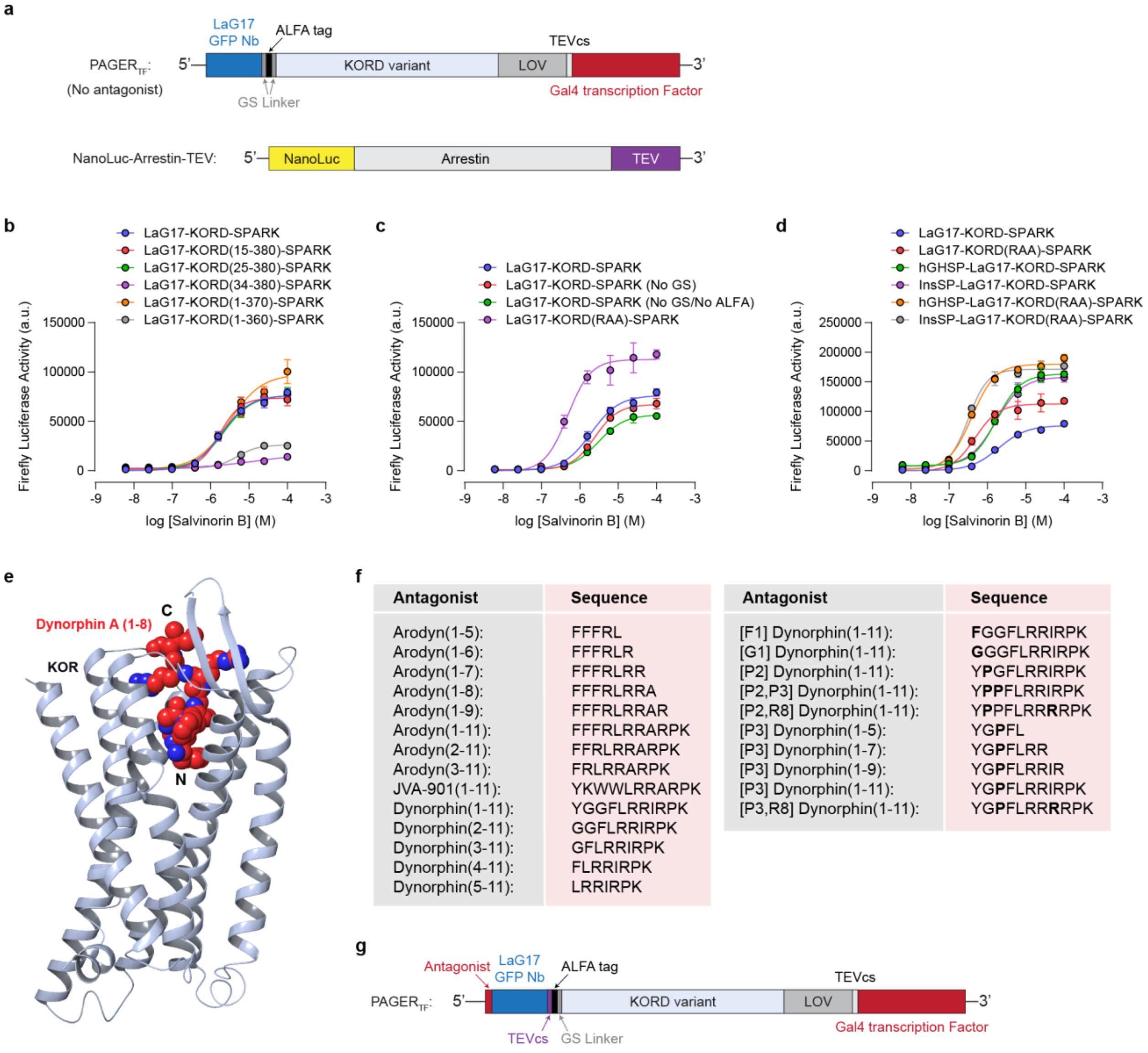


**Extended Data Figure 1. Construct design and optimization of PAGER$_{TF}$. a**, Domain structures of PAGER$_{TF}$ (without antagonist) and Arrestin-TEV constructs. **b-d**, Optimization of the PAGER$_{TF}$ construct. SalB dose response curves of PAGER$_{TF}$ constructs with various truncations (**b**), linkers or mutation (**c**), and signal peptides (**d**). Full length KORD (V360A/R361A) with IL-2 signal peptide was chosen as the optimal PAGER$_{TF}$ construct and was used in all subsequent experiments. Data representative of $n = 2$ independent experiments. **e,** Crystal structure of the Kappa-opioid receptor (KOR) in complex with dynorphin A (PDB: 8F7W). Dynorphin binds KOR with its N-terminus buried in the orthosteric binding pocket. **f**, List of candidate peptide antagonists screened in PAGER$_{TF}$. **g**, Domain structure of PAGER$_{TF}$ including an N-terminal antagonist and TEVcs in the extracellular linker.

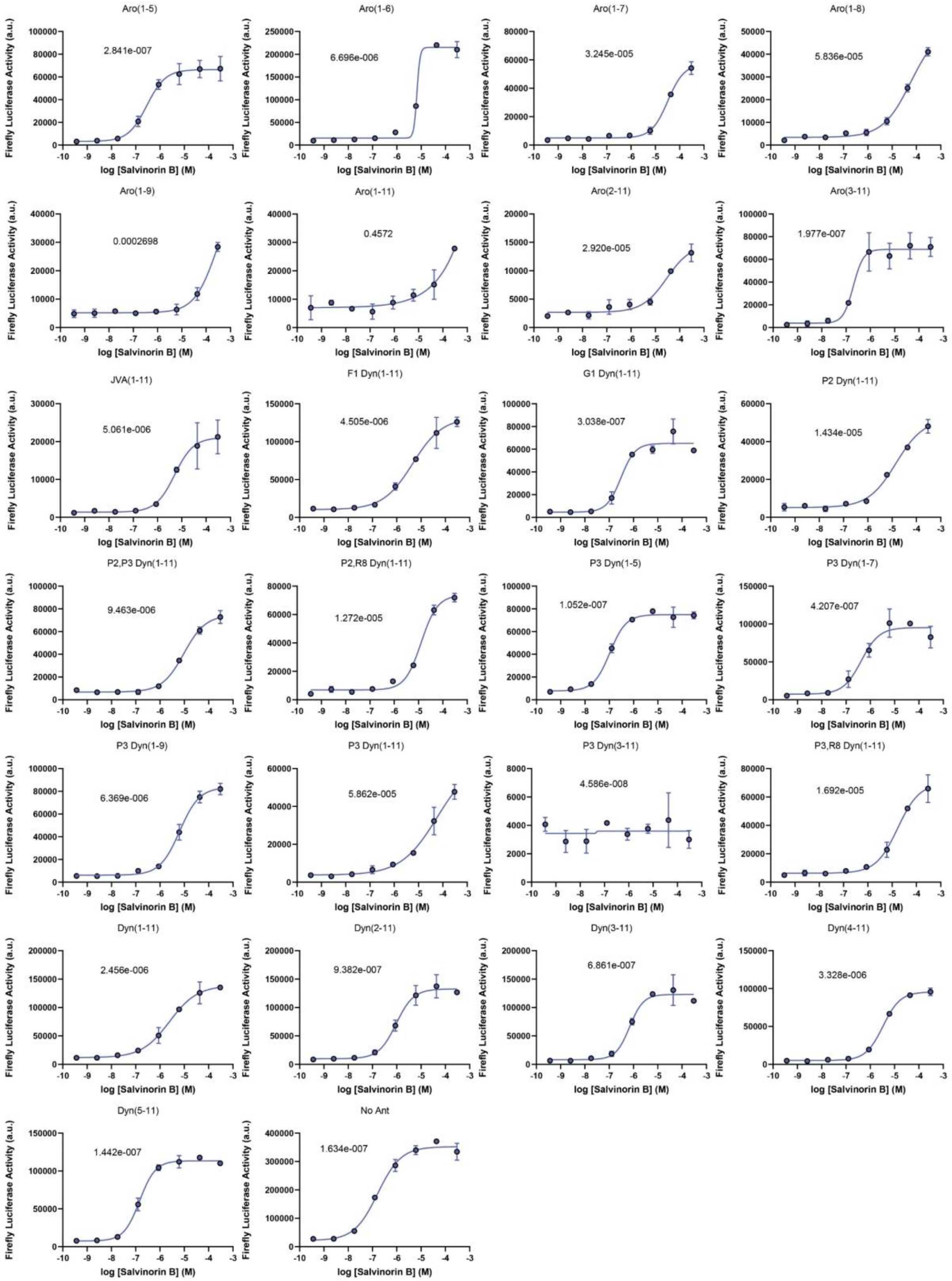


**Extended Data Figure 2. Screening candidate peptide antagonists in PAGER$_{TF}$.** SalB dose response curves for PAGER$_{TF}$ constructs with various candidate peptide antagonists. Sequences of candidate peptide antagonists are included in **Extended Data Fig. 1f**. Peptides with antagonistic activity shift the SalB response curve and EC50 to the right relative to no antagonist. EC50s are plotted in **Fig. 1d**. Data representative of $n = 2$ independent experiment.

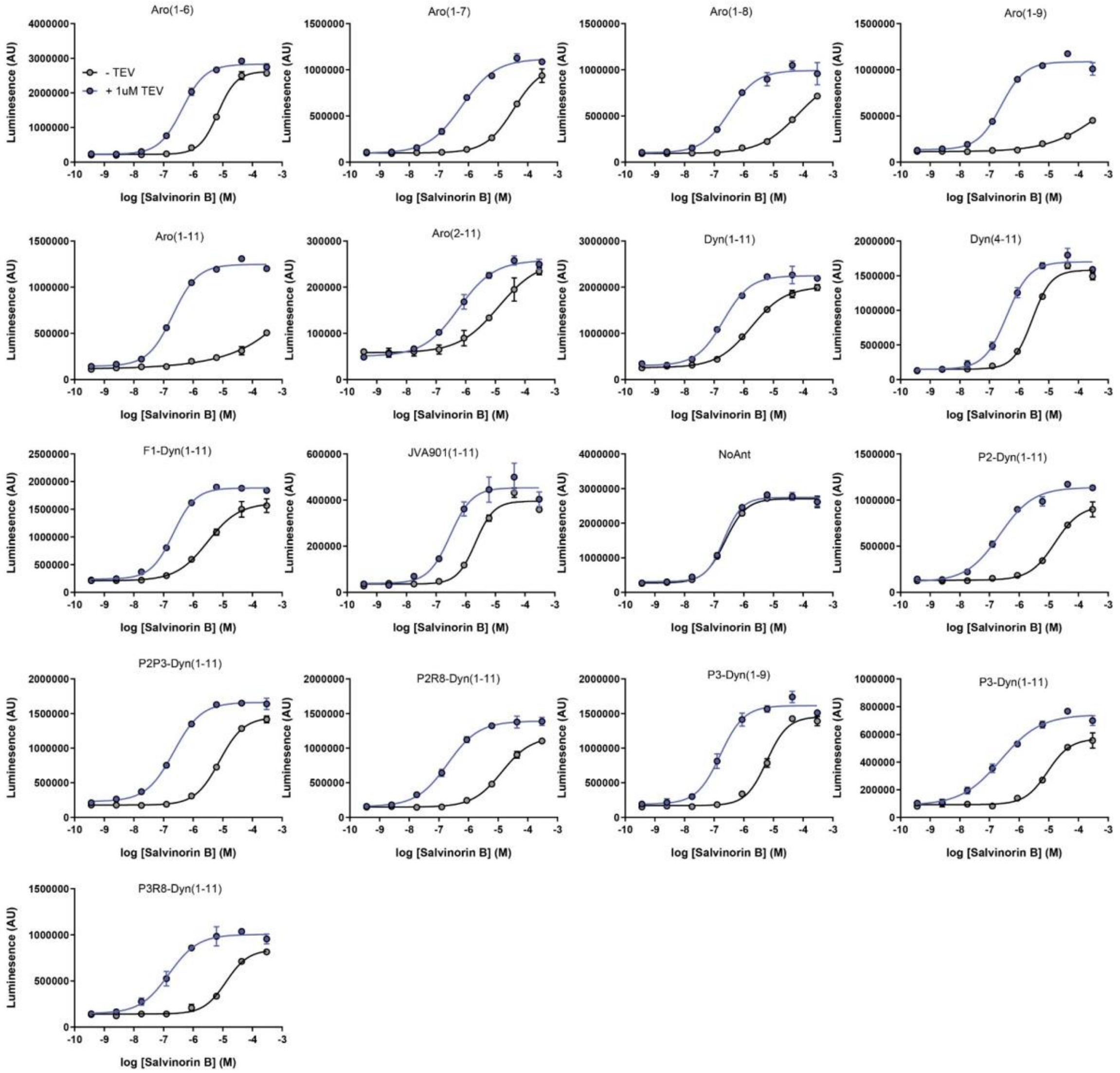


**Extended Data Figure 3. Screening PAGER$_{TF}$ constructs for reversible auto-inhibition using recombinant TEV protease.** SalB dose response curves for PAGER$_{TF}$ constructs with various candidate peptide antagonists with and without 90 min pre-incubation with 1 μM TEV protease. Constructs that display reversible auto-inhibition have left-shifted SalB dose response curves and EC50s when pre-incubation with TEV protease is included. The four constructs selected for further screening are indicated with red boxes. Data for these constructs are also included in **Fig. 1f-g**. Data representative of $n = 2$ independent experiment.

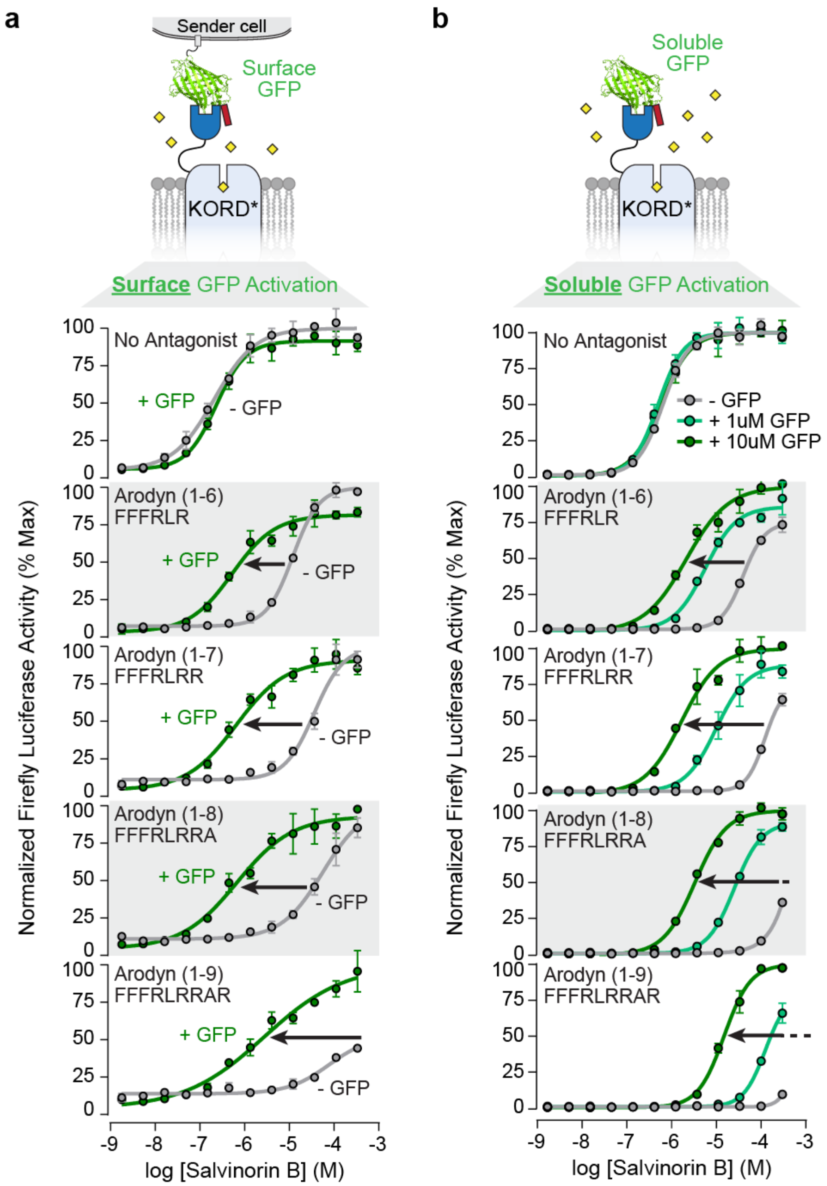


**Extended Data Figure 4. Testing selected PAGER$_{TF}$ constructs for response to surface or soluble GFP antigen. a**, SalB dose response curves for various PAGER$_{TF}$ constructs with or without co-culture with surface GFP-expressing sender cells. **b**, SalB dose response curves for various PAGER$_{TF}$ constructs with or without soluble GFP. Data presented here is an extended version of data included in Fig. 1j and 1m. Data representative of *n* = 1 independent experiment.

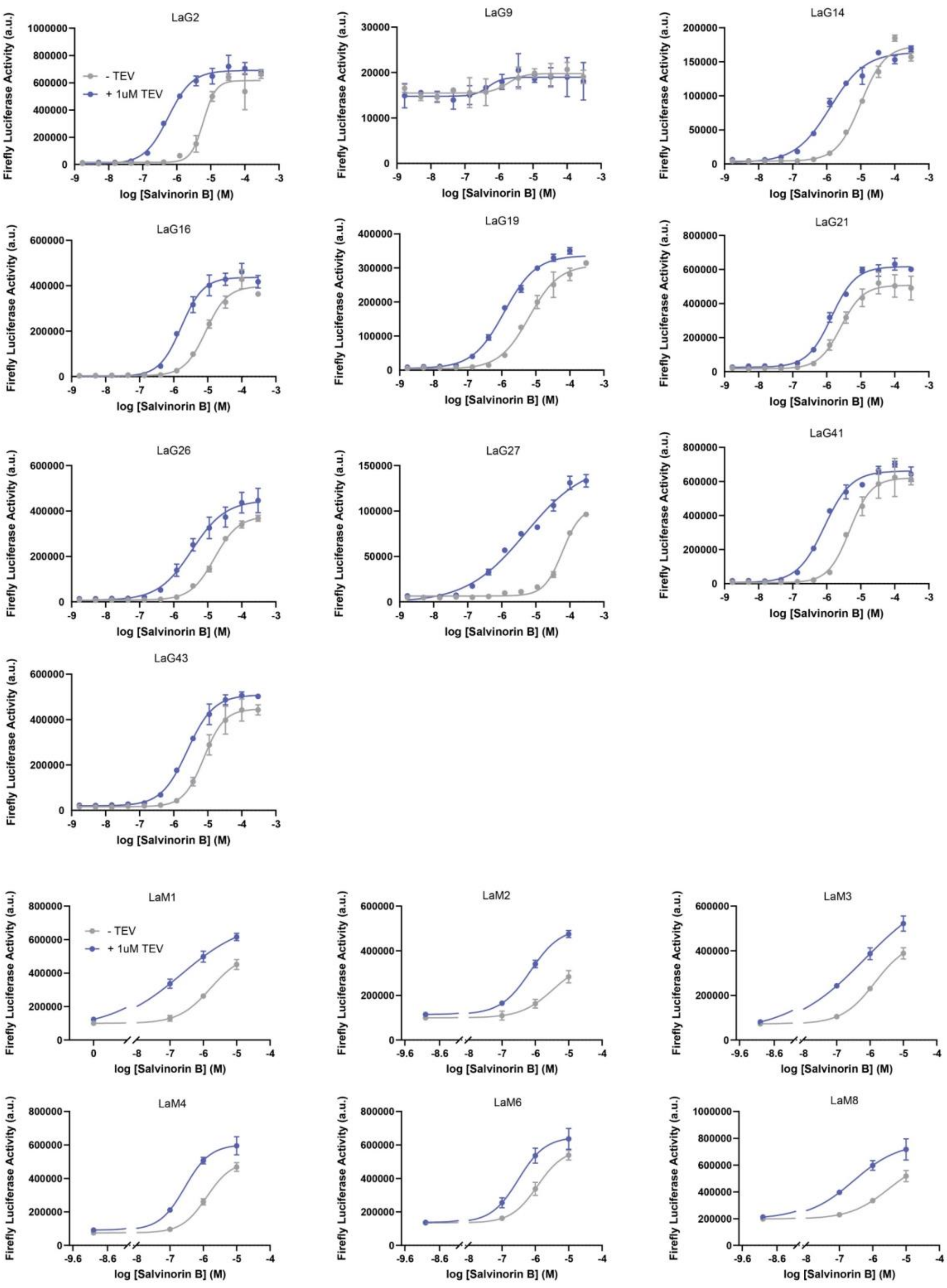


**Extended Data Figure 5. Testing α-GFP and α-mCherry PAGER$_{TF}$ constructs for reversible auto-inhibition using recombinant TEV protease.** SalB dose response curves for α-GFP and α-mCherry PAGER$_{TF}$ constructs containing the indicated GFP and mCherry specific nanobodies, with and without 90 min pre-incubation with 1 μM TEV protease. Data representative of *n* = 1 independent experiment.

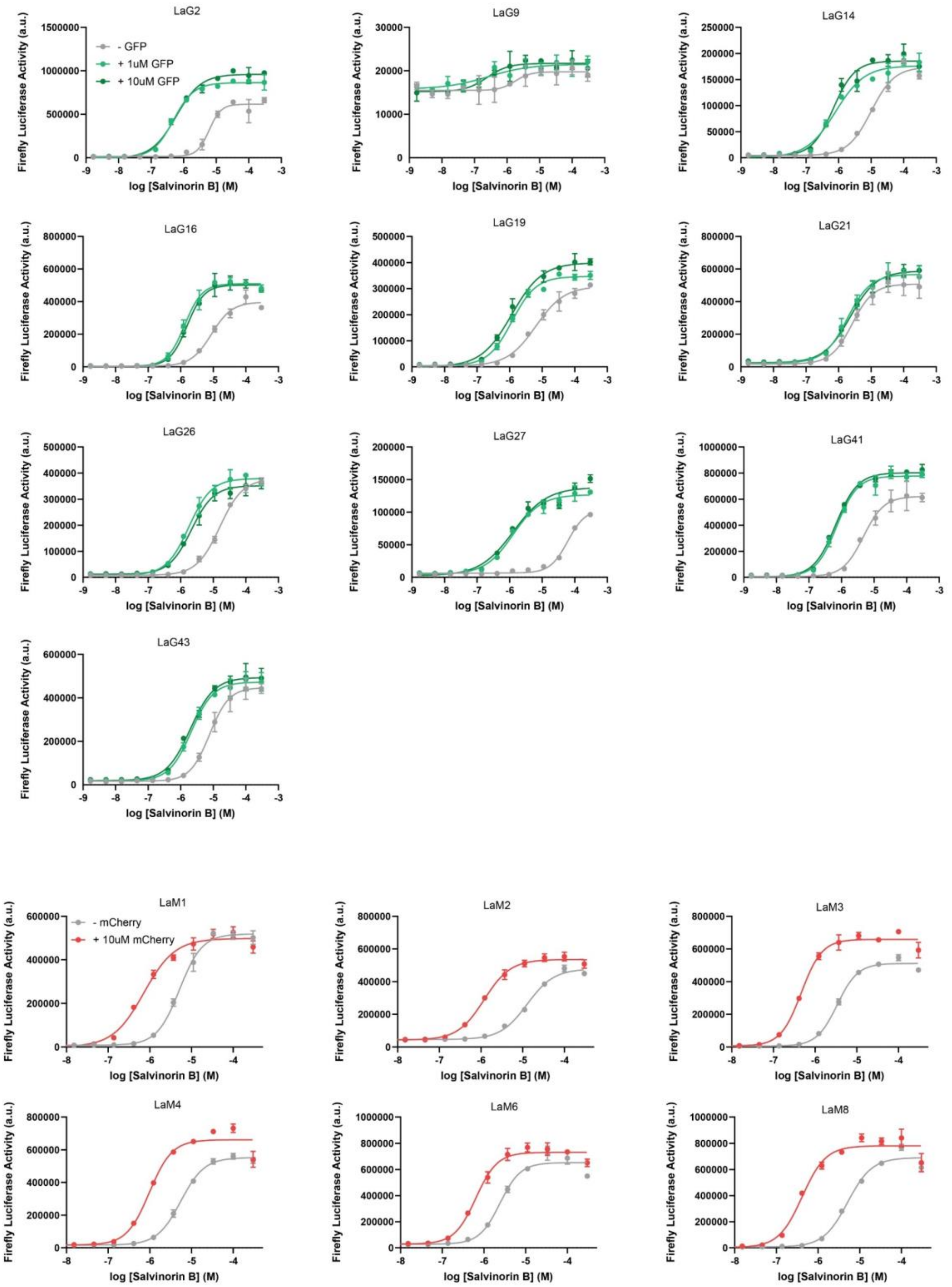


**Extended Data Figure 6. Testing α-GFP and α-mCherry PAGER$_{TF}$ constructs for response to soluble cognate antigen.** SalB dose response curves for α-GFP and α-mCherry PAGER$_{TF}$ constructs with or without soluble cognate antigen. Data representative of $n = 1$ independent experiment.

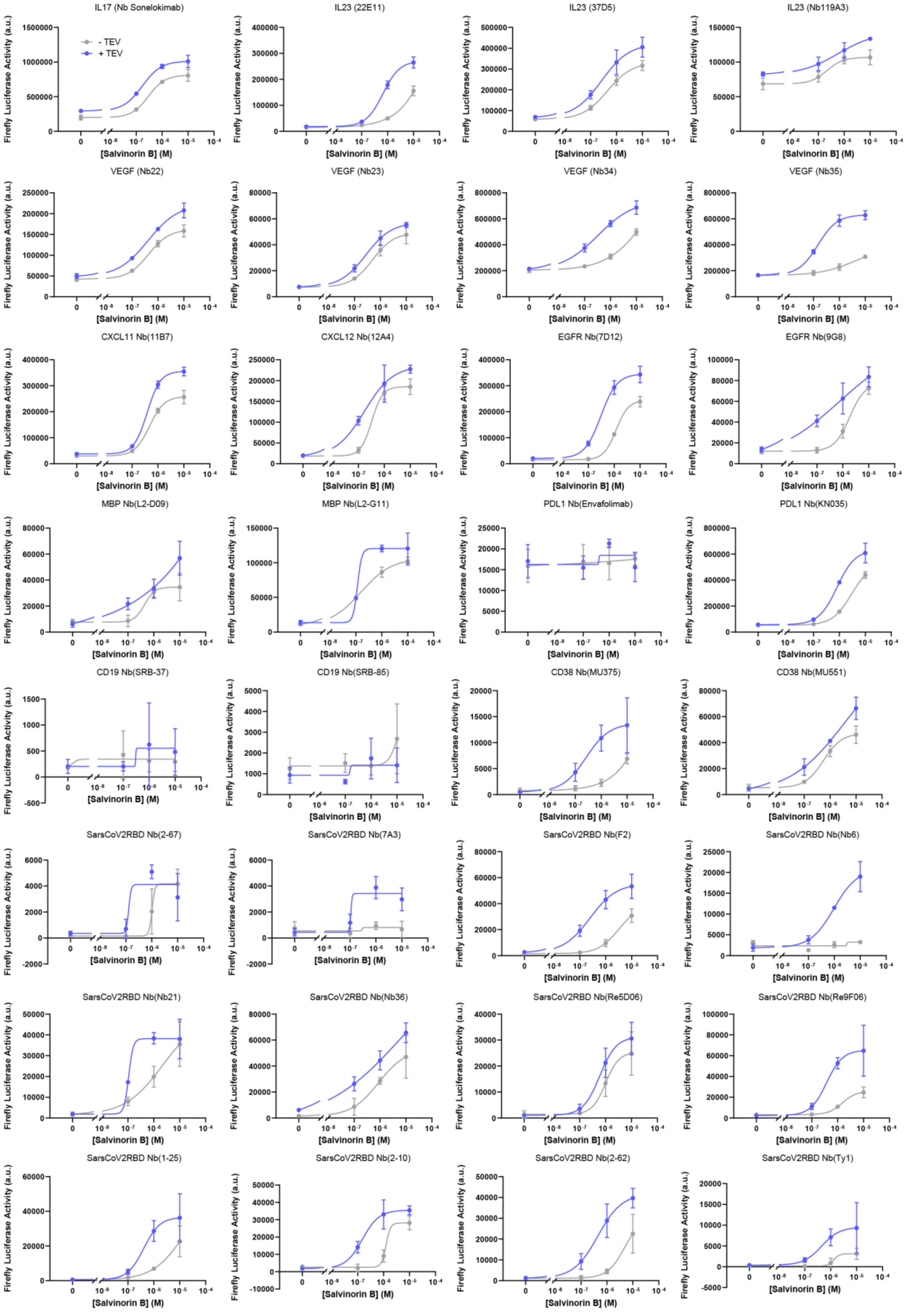
IL17 (Nb Sonelokimab)
- TEV
+ TEV
IL23 (22E11)
IL23 (37D5)
IL23 (Nb119A3)
VEGF (Nb22)
VEGF (Nb23)
VEGF (Nb34)
VEGF (Nb35)
CXCL11 Nb(11B7)
CXCL12 Nb(12A4)
EGFR Nb(7D12)
EGFR Nb(9G8)
MBP Nb(L2-D09)
MBP Nb(L2-G11)
PDL1 Nb(Envafolimab)
PDL1 Nb(KN035)
CD19 Nb(SRB-37)
CD19 Nb(SRB-85)
CD38 Nb(MU375)
CD38 Nb(MU551)
SarsCoV2RBD Nb(2-67)
SarsCoV2RBD Nb(7A3)
SarsCoV2RBD Nb(F2)
SarsCoV2RBD Nb(Nb6)
SarsCoV2RBD Nb(Nb21)
SarsCoV2RBD Nb(Nb36)
SarsCoV2RBD Nb(Re5D06)
SarsCoV2RBD Nb(Re9F06)
SarsCoV2RBD Nb(1-25)
SarsCoV2RBD Nb(2-10)
SarsCoV2RBD Nb(2-62)
SarsCoV2RBD Nb(Ty1)
Firefly Luciferase Activity (a.u.)
[Salvinorin B] (M)

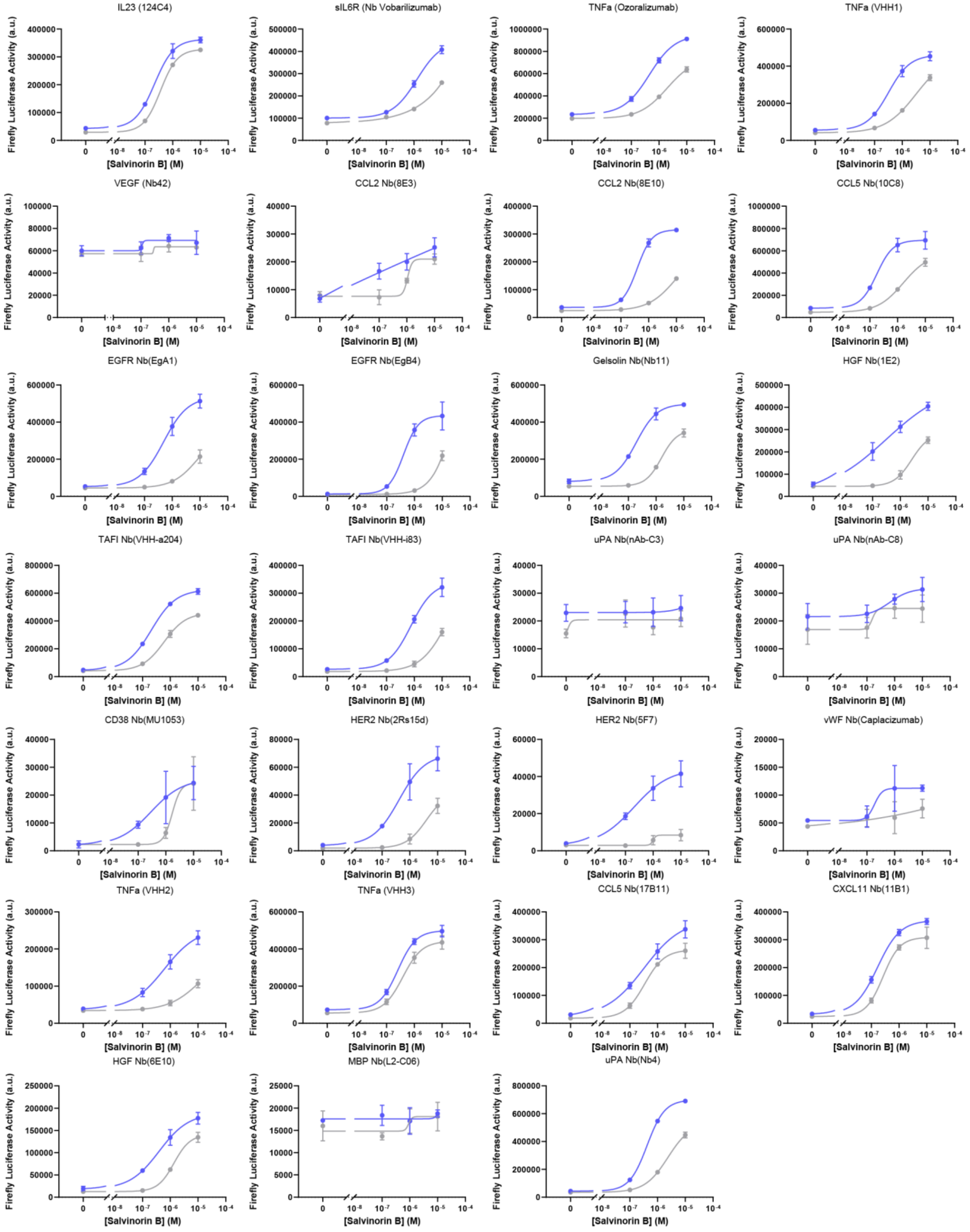


**Extended Data Figure 7. Testing PAGER$_{TF}$ constructs with various nanobodies for reversible auto-inhibition using recombinant TEV protease.** SalB dose response curves for PAGER$_{TF}$ constructs containing the indicated nanobodies, with and without 90 min pre-incubation with 1 μM TEV protease. Data representative of $n = 1$ independent experiment.

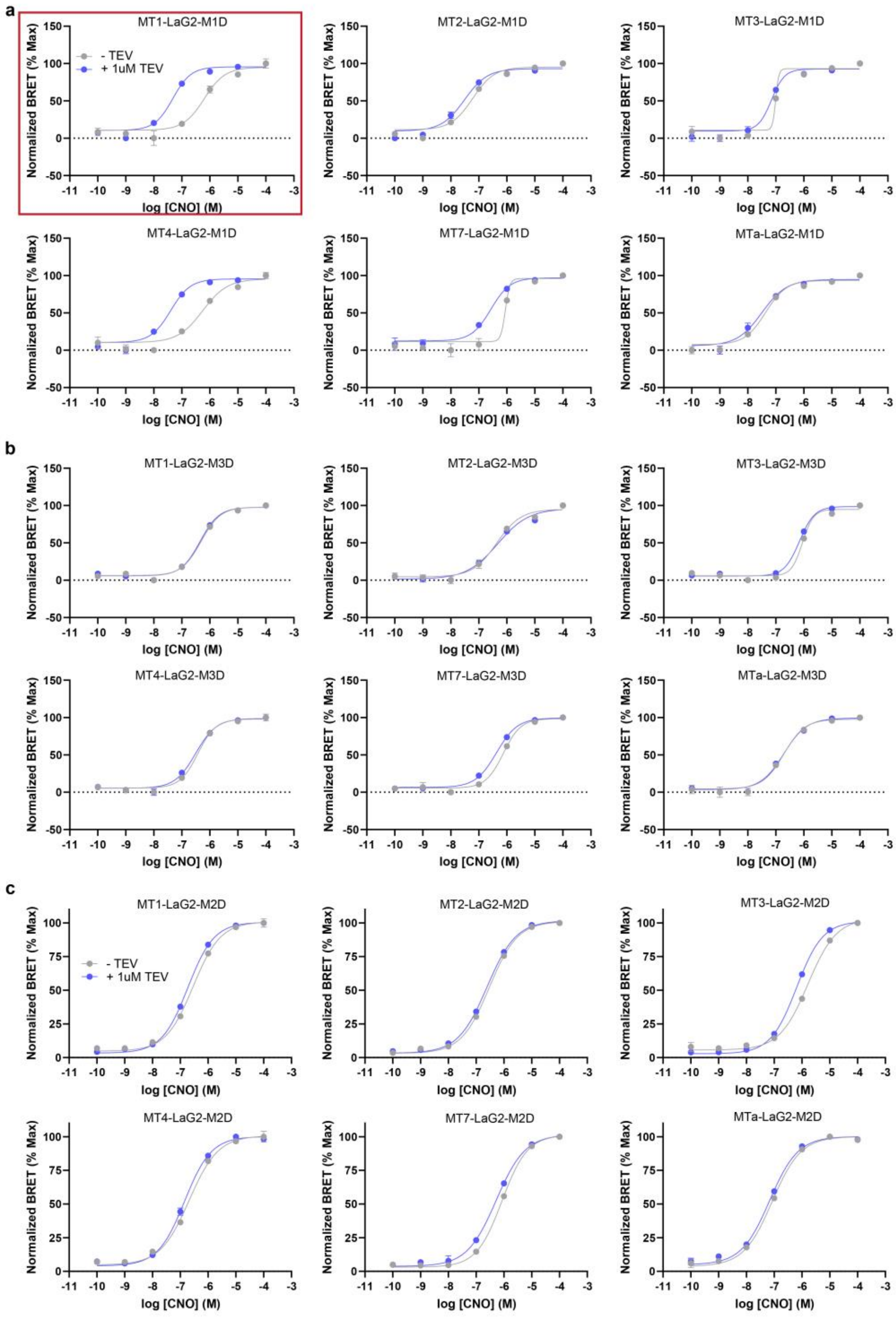

a
MT1-LaG2-M1D
- TEV
+ 1uM TEV
MT2-LaG2-M1D
MT3-LaG2-M1D
MT4-LaG2-M1D
MT7-LaG2-M1D
MTa-LaG2-M1D
b
MT1-LaG2-M3D
MT2-LaG2-M3D
MT3-LaG2-M3D
MT4-LaG2-M3D
MT7-LaG2-M3D
MTa-LaG2-M3D
c
MT1-LaG2-M2D
- TEV
+ 1uM TEV
MT2-LaG2-M2D
MT3-LaG2-M2D
MT4-LaG2-M2D
MT7-LaG2-M2D
MTa-LaG2-M2D
Normalized BRET (% Max)
log [CNO] (M)

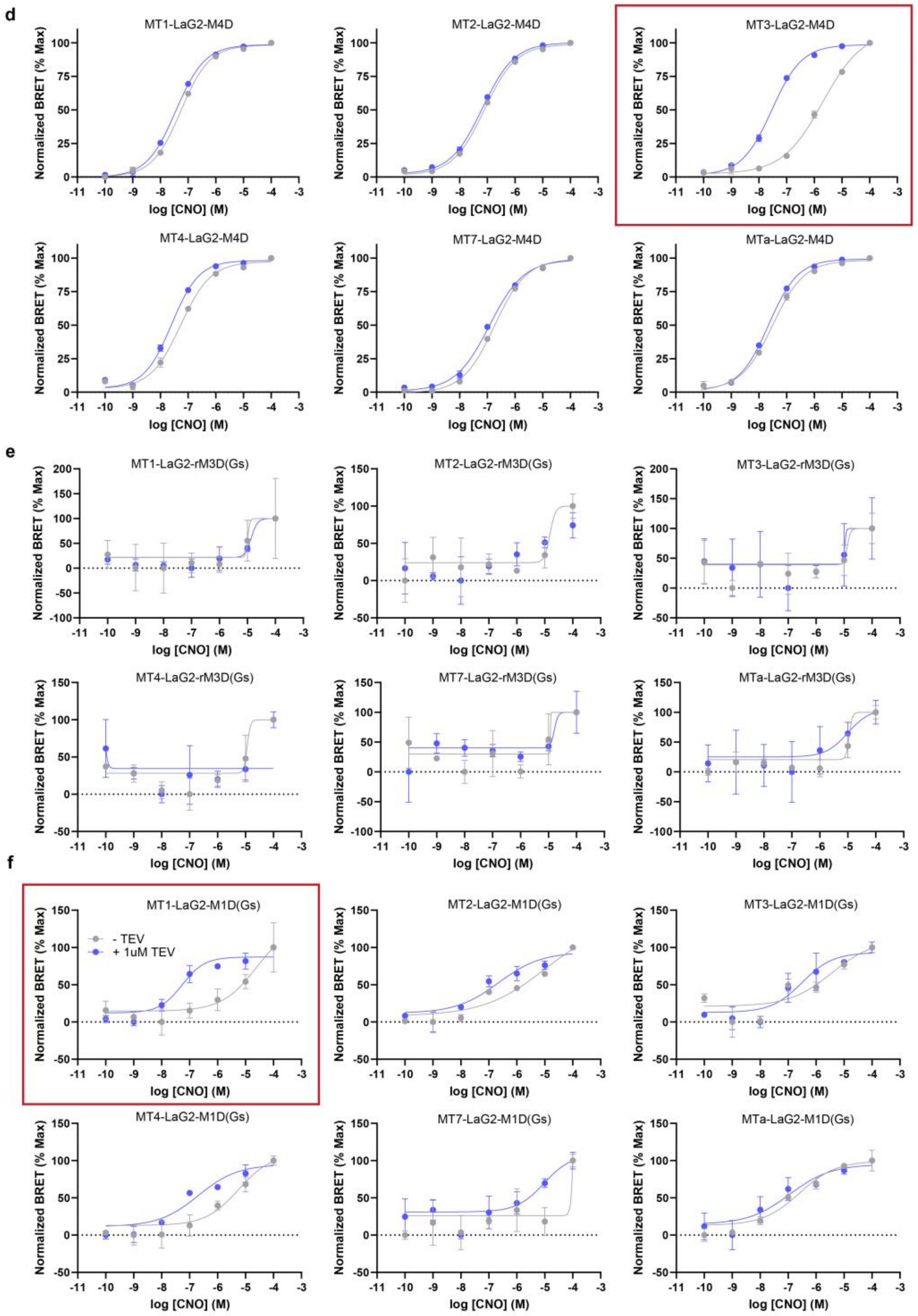


**Extended Data Figure 8. Screening muscarinic toxins in PAGER$_G$ constructs for reversible auto-inhibition using recombinant TEV protease a-f,** SalB dose-responsive curves for muscarinic toxin screening in hM1D-based PAGER (**a**, Fig. 3b), hM3Dq-baesd PAGER (**b**, Extended Data Fig. 6g)**,** hM2Di-based PAGER (**c**, Extended Data Fig. 6i), hM4D-based PAGER(**d**, Fig. 3d), rM3D-based PAGER (**e**, Extended Data fig. 6h)**, and chimeric hM1Ds-based PAGER(f**, Fig. 3c). Data representative of *n* = 1 independent experiment.

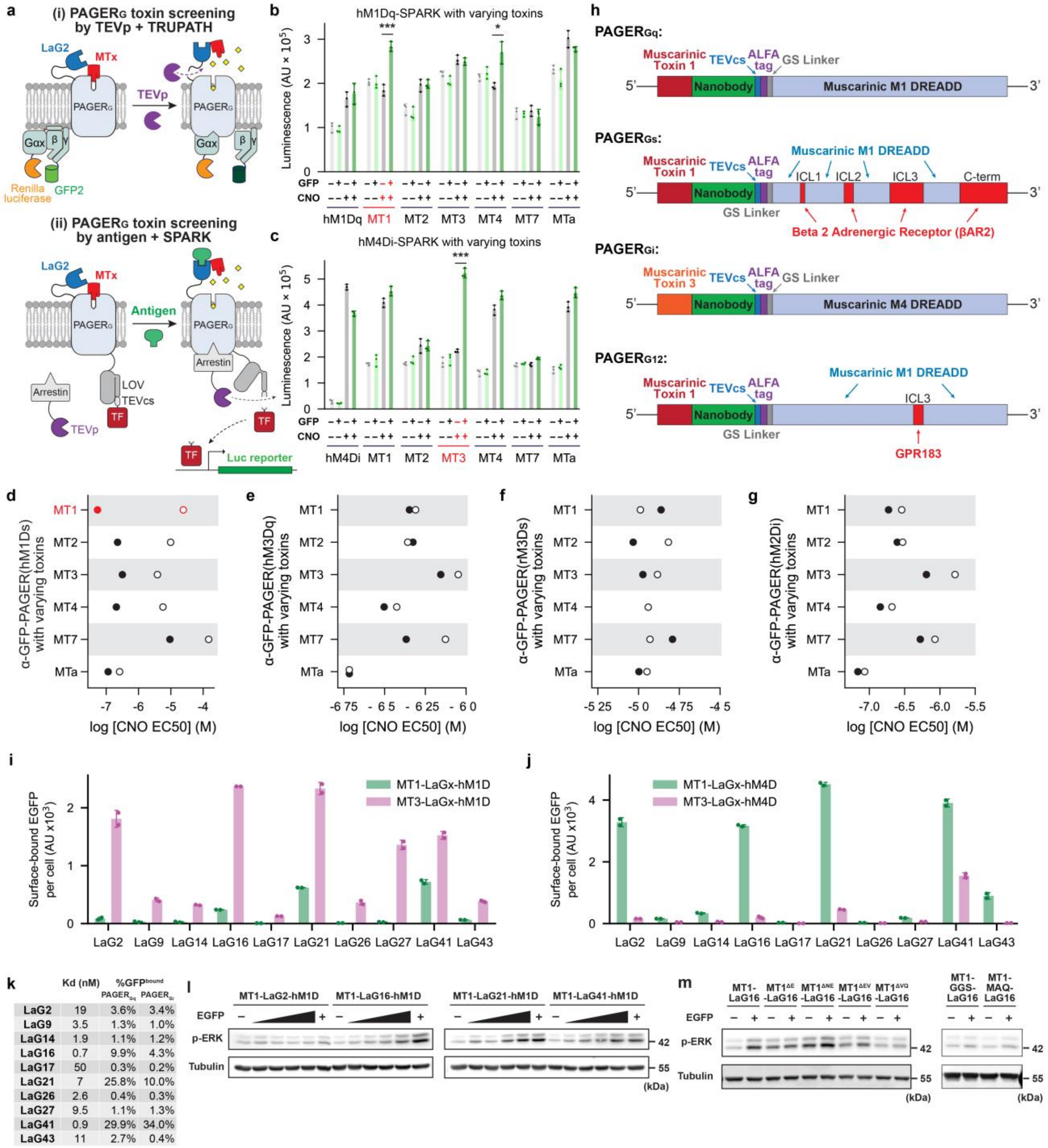


| | Kd (nM) | %GFP^bound PAGER$_{Gq}$ | %GFP^bound PAGER$_{Gi}$ |
|---|---|---|---|
| LaG2 | 19 | 3.6% | 3.4% |
| LaG9 | 3.5 | 1.3% | 1.0% |
| LaG14 | 1.9 | 1.1% | 1.2% |
| LaG16 | 0.7 | 9.9% | 4.3% |
| LaG17 | 50 | 0.3% | 0.2% |
| LaG21 | 7 | 25.8% | 10.0% |
| LaG26 | 2.6 | 0.4% | 0.3% |
| LaG27 | 9.5 | 1.1% | 1.3% |
| LaG41 | 0.9 | 29.9% | 34.0% |
| LaG43 | 11 | 2.7% | 0.4% |

**Extended Data Figure 9. Development and characterization of PAGER$_G$. a,** Schematics of TRUPATH BRET assay (i) SPARK assay (ii) used to measure G protein activation in response to G-coupled PAGERs for screening muscarinic toxins (MTs). **b,** Screening toxin variants for auto-inhibition of hM1Dq-based PAGER-TF. HEK cells expressing the indicated variant were treated with 1 uM GFP and 10 µM CNO for 15 minutes. 8 hours later, luciferase reporter expression was quantified. MT1 toxin was best for gating PAGER$_{Gq}$ consistent with the screening based on TRUPATH assay. **c,** Same assay as (c) for hM4Di-based PAGER-TF. MT3 toxin was best for gating PAGER$_{Gi}$, consistent with the screening based on TRUPATH assay. **d,** Screening toxin variants for auto-inhibition of rM1Ds (where ICL2–3 in rM3Ds were grafted into hM1Dq). HEK cells expressing the indicated variant were stimulated with varying concentrations of CNO, with (filled) or without (open) prior TEVp treatment to cleave off the toxin. $PAGER_{Gs}$ activation was measured with TRUPATH BRET assay. Full drug response curves in Extended Data Fig. 7. MT1 toxin was best for gating rM1Ds. **e–g,** Same assay as in (d) but for hM3Dq, rM3Ds, and hM2Di. hM3Dq, rM3Ds, and hM2Di could not be gated by any of the toxins tested. The only MTx with reported high affinity for M3 is MTα, but it failed to show inhibition against hM3Dq nor rM3Ds. Data in d-g are representative of $n = 1$ independent experiment. **h,** Domain structures of optimized $PAGER_{Gq}$, $PAGER_{Gs}$, $PAGER_{Gi}$, and $PAGER_{G12}$. **i,** Screening GFP nanobodies in α-GFP-$PAGER_{Gq}$ (based on hM1Dq) with either MT1 (green) or MT3 (magenta) as autoinhibitory domain. MT1 has higher reported affinity to M1 receptor than MT3. **j,** Same assay as (j) but with α-GFP-$PAGER_{Gi}$ (based on hM4Di). MT3 has higher reported affinity to M4 receptor than MT1.**k,** A table summarizing reported affinity ($K_d$) of α-GFP nanobodies along with the amount of GFP bound onto $PAGER_{Gx}$ (divided by the maximum amount of GFP bound in the lack of any antagonism). **l,** Western blots showing phosphorylation of endogenous ERK in response to GFP and CNO activation of $PAGER_{Gq}$ with four different GFP nanobodies. HEK cells expressing indicated α-GFP-PAGER was stimulated with 100 nM CNO and 0, 0.5, 5, 50, 500, or 5000 μM antigen for 3 min, before cell lysis and analysis. Phospho-ERK increase started to be apparent with 50 nM antigen. **m,** Similar assay as (l) with 1 μM antigen but with $PAGER_{Gq}$ with either truncation or extension between MT1 and nanobody.

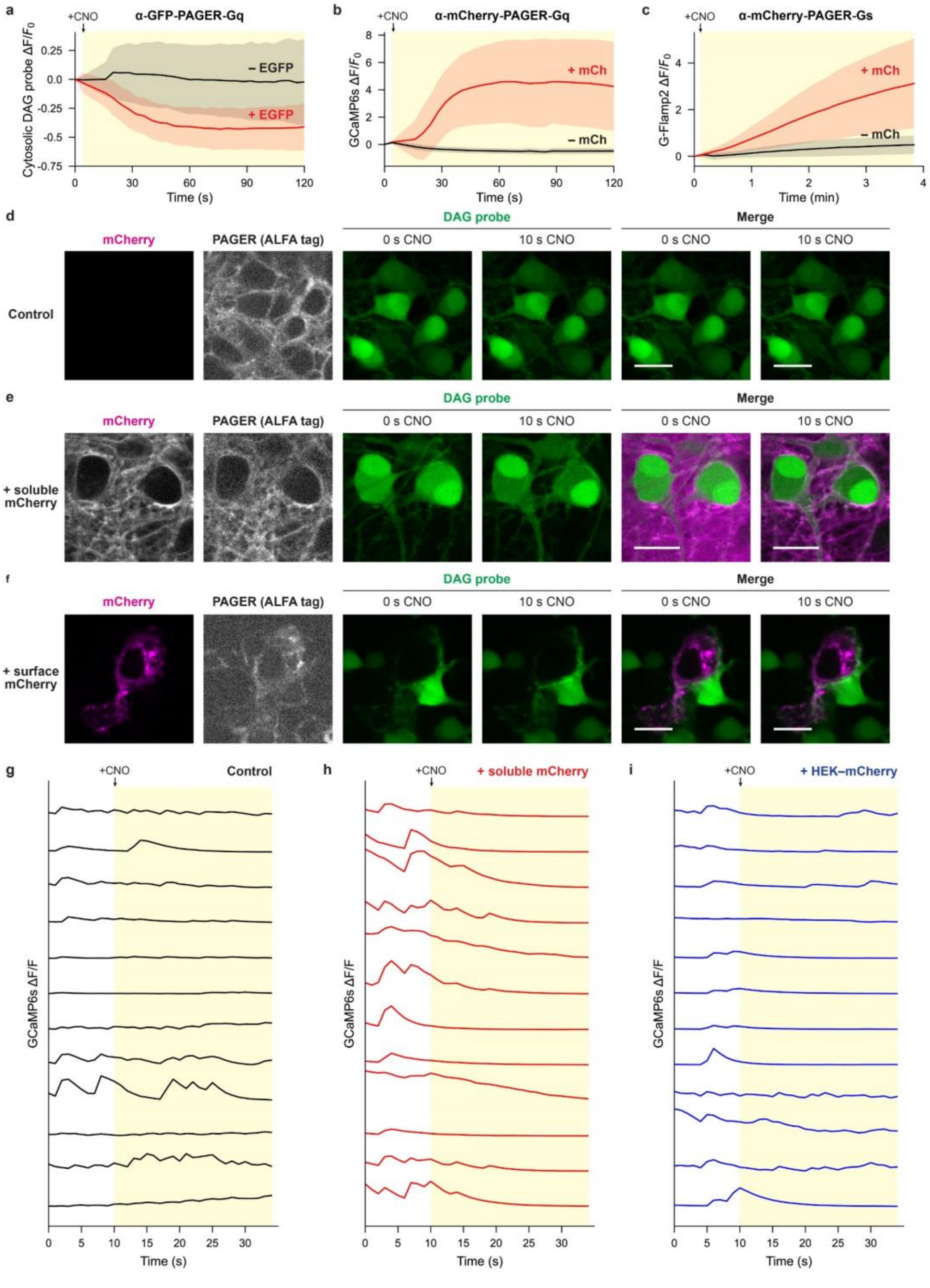


**Extended Data Figure 10. PAGER$_G$ mediates antigen-dependent control of cell signaling and behavior. a–c,** Time-course plots for data in Fig. 4b–c (a), Fig. 4d–e (b), Fig. 4f–g (c). CNO was added at t=4s to the final concentration of 100 nM. **d–f,** More representative images for the assay described in Fig. 4k–l. Neurons expressing α-mCherry-PAGER-Gq and DAG probe were stimulated with 100 nM CNO with no mCherry (d), 1 μM soluble mCherry (e), and surface mCherry (f) and imaged over time. **g–i,** Individual calcium traces for the assay described in Fig. 4m–o. Rat cortical neurons co-expressing α-mCherry-PAGER-Gi and GCaMP6s, with no antigen (g), with 1 μM mCherry (h), or co-plated with HEK cells expressing surface mCherry (i). CNO was added at t=10 s to the final concentration of 30 nM.

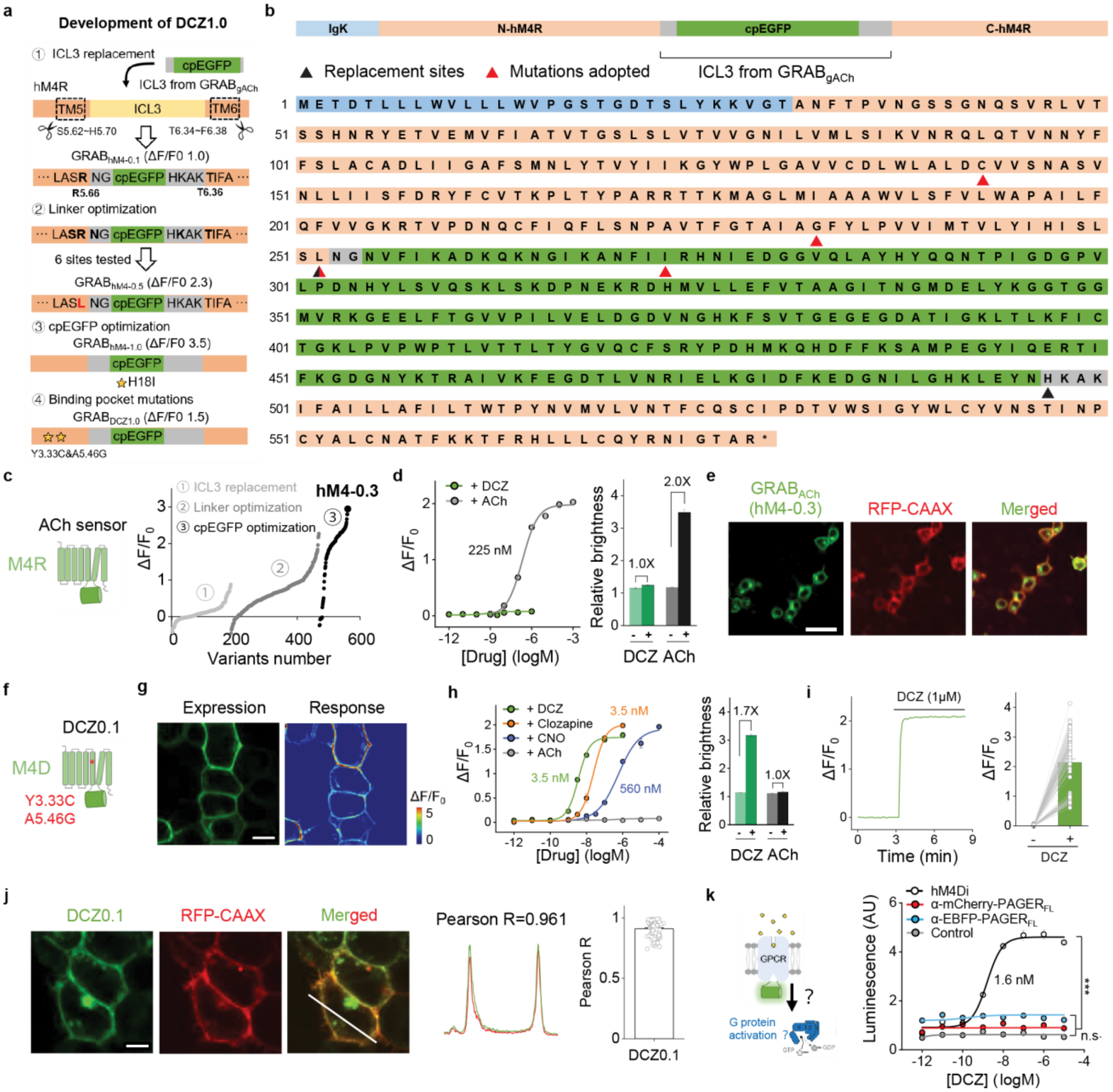

**Extended Data Fig. 11. Engineering PAGER$_{FL}$.** **a**, Summary of how GRAB$_{DCZ1.0}$ was developed. GRAB$_{DCZ1.0}$ is the receptor portion of PAGER$_{FL}$, lacking the nanobody and toxin. Starting from M4R, we ① screened cpEGFP insertion sites within the ICL3 loop in gACh sensor[2]; ② optimized key residues (shown in bold) in the linkers; and ③ optimized critical residues in cpEGFP. **b**, Sequence of optimized GRAB$_{DCZ1.0}$ sensor. The residues related to ICL3 replacement and mutations incorporated during the optimization process are marked. **c**, Summary of GRAB$_{ACh}$ screening and optimization. The number of GRAB$_{ACh}$ variants tested during each optimization process is shown in x-axis. The final optimized variant was named hM4-1.0. **d**, Response (left) and relative brightness (right) of GRAB$_{ACh}$ (hM4-1.0) to various concentrations of ACh and DCZ. n = 3 wells containing 100 to 300 cells per well. **e**, Membrane expression of GRAB$_{ACh}$ in HEK293T cells. RFP-CAAX is a membrane-targeted RFP. Scale bars, 50 µm. **f**, GRAB$_{DCZ1.0}$ has two additional mutations compared to GRAB$_{ACh}$. **g**, Representative images of expression and response of GRAB$_{DCZ1.0}$ sensor to 1 µM DCZ. Scale bars, 10 µm. **h**, Drug response curves (left) and relative brightness (right) of GRAB$_{DCZ1.0.}$ n = 3 wells containing 100 to 300 cells per well. **i**, Fluorescence time traces (left) showing the rate of EGFP fluorescence onset after 1 μM DCZ addition. Group summary (right) of $\Delta F/F_0$ with or without DCZ addition. n = 126 cells from 3 coverslips. **j**, Example fluorescence images and intensity line scan profiles of $GRAB_{DCZ1.0}$ (green), RFP-CAAX (red), and merged image in the presence of DCZ (1 μM). The white line indicated the ROI for intensity profiling, and the Pearson R was calculated and used to indicate the membrane trafficking index of the sensor. n = 112 cells from 3 coverslips. Scale bars, 10 μm. **k**, G protein and coupling were measured using the split-luciferase complementation assay[3] in cells expressing αmCherry-$PAGER_{FL}$ (red), αEBFP-$PAGER_{FL}$ (blue) in the presence of the indicated concentrations of DCZ; The DCZ-responsive DREADD hM4Di (black) was used as a positive control and no receptor (grey) was used as a negative control. n = 3 wells each. AU, arbitrary units.
